# Medical data and machine learning improve power of stroke genome-wide association studies

**DOI:** 10.1101/2020.01.22.915397

**Authors:** Phyllis M. Thangaraj, Undina Gisladottir, Nicholas P. Tatonetti

## Abstract

Genome-wide association studies (GWAS) may require enrollment of up to millions of participants to power variant discovery. This requires manual curation of cases and controls with large-scale collaborations. Biobanks connected to electronic health records (EHR) can facilitate these studies by using data from clinical care systems, like billing diagnosis codes, as phenotypes. These systems, however, do not define adjudicated cases and controls. We developed QTPhenProxy, a machine learning model that adds nuance to cohort classification by assigning everyone in a cohort a probability of having the study disease. We then ran a GWAS using the probabilities as a quantitative trait. With an order of magnitude fewer cases than the largest stroke GWAS, our method outperformed previous methods at replicating known variants in stroke and discovered a novel variant in *ABCG8* associated with intracerebral hemorrhage in the UK Biobank that replicated in the MEGASTROKE GWA meta-analysis. QTPhenProxy expands traditional phenotyping to improve the power of GWAS.

## 1. Introduction

Large genetic repositories connected to electronic health records (EHR) promise the ability to perform thousands of genetic studies using data routinely captured in clinical practice. High-throughput identification of cases and controls can be difficult, however, due to time-consuming chart review and incompleteness of medical records. Current disease genome-wide association studies develop case-control sets in which power relies on a large amount of pure cases, and the missingness of EHR data could prevent some cases from discovery in a high-throughput manner (McCarthy et al., 2008; Weiskopf et al., 2013; DeBoever et al., 2019). In addition, extreme case-control imbalance in biobanks can lead to increased type 1 error when running linear mixed model genome-wide association analysis (Zhou et al., 2018). An incorporation of additional accessible EHR data could improve case curation sensitivity. In addition, many diseases such as stroke result from a combination of gene and environmental interactions, and there is significant overlap with co-morbidities in genome-wide significant variants (Malik et al., 2018a). Therefore, it is difficult to confirm every person without the event a control, suggesting the utility of a disease likelihood assignment (McCarthy et al., 2008; Yang et al., 2009).

The definition of the disease phenotype influences the success of detecting a genetic signal since power is generally calculated by number of cases and controls (Maher, 2008; Zaykin and Zhivotovsky, 2005). We propose that including EHR information about comorbidities (Kohane, 2011) and other health information about our subject to trait assignment can improve the power of GWAS. Past studies have shown that incorporating diagnosis count improved the power of genetic studies, and the addition of patient questionnaires and genetic correlations to hospital records improved detection of cases for genome-wide association studies (Sinnott et al., 2018; DeBoever et al., 2019; Liu et al., 2017; Ruderfer et al., 2019). Incorporating EHR data to develop a probability of suicide attempt also improved the power of its genome-wide association study (Ruderfer et al., 2019). We argue that including several modalities of health data to estimate assignment of case probability can improve the power of genomic studies. For example, the most successful genome-wide association study for stroke required 40,585 cases and 406,111 controls (Malik et al., 2018a). We hypothesize that we can discover genome-wide significant variants associated with stroke with a fraction of those cases (4,354) by incorporating EHR information of 337,147 participants in the UK Biobank Cohort into a stroke phenotype score to be used as quantitative trait.

In this study, we use machine learning methods to expand sample cohorts by assigning every patient a probability of disease. We hypothesize that the output of a supervised machine learning classifier, trained on the EHR data of a small number of known cases and controls, can be used as a proxy variable for stroke and will be an efficient strategy for expanding cohort size (See Graphical Abstract). We demonstrate that our quantitative proxy trait can improve power over its respective binary trait in ischemic stroke. We also show that the new variants discovered are known in similar cardiovascular and neurological diseases. We find up to 7 LD independent loci that pass genome-wide significance and conditional analysis, and the majority of the associated genes are known to be associated with stroke or cardiovascular disease. For stroke and its subtypes ischemic stroke, subarachnoid hemorrhage, and intracerebral hemorrhage, QTPhenProxy recovered known and discovered new stroke variants, such as in *ABCG8*, with an order of magnitude fewer cases than traditional genome-wide association studies.

## 2. Methods

### 2.1. QTPhenProxy Phenotyping Model

We gathered clinical features of primary ICD10 dignosis codes, OCPS4 procedure codes (UK Biobank code 42100), medications (UK Biobank code 20003), race/ethnicity (UK Biobank code 1001), and age. 2018 served as the age end point. We mapped the OCPS4 procedure codes (UK Biobank code 240) to SNOMED-CT codes and the UK Biobank medication codes to RxNorm codes by name (National Library of Medicine, 2019b,a). We also gathered data of self reported and first occurrence of all stroke, ischemic stroke, sub-arachnoid stroke, and intracerebral hemorrhage. We made a large matrix in which each extracted EHR feature was a binary variable based on the presence or absence of the feature. We dichotomized age as greater than or equal or less than 50 years. For each disease, we then defined the cases from the ICD10 code combinations described in the UK Biobank’s phenotyping algorithm (Schnier et al., 2017). We then trained 5 different classifiers: 1) logistic regression with elastic net penalty, 2) logistic regression with L1 penalty, 3) random forest, 4) adaboost, 5) gradient boosting classifiers on 50% of the cases and an equal number of controls. Controls were identified as subjects without any ICD10 codes within the same category as the Clinical Modification Clinical Classifications Software tool (Healthcare Cost and Utilization Project). We then applied the trained algorithm to the whole UK Biobank, resulting in a model probability or quantitative trait proxy for each subject and each disease. For comparison, another phenotype file with binary assignment of case and control for each disease was prepared as well. Table 1 shows the number of cases available for each disease.

**Table 1:** Number of genome-wide significant loci and proportion of known stroke variants that reach nominal significance for each model. *IscStroke*: Ischemic Stroke, *SAHStroke*: Subarachnoid Hemorrhage, *ICHStroke*: Intracerebral Hemorrhage, *N Cases*: number of cases, *hits*: Number of genome-wide significant loci, sens: sensitivity measure for each model GWAS and binary trait GWAS, measures what proportion of known stroke variants reach genome-wide significance in each association test. *RF*: Random Forest model, *EN*: Logistic Regression with elastic net penalty model.

| Disease | N Cases | Binary Hits | Bin Sens | RF Hits | RF Sens | EN Hits | EN Sens |
| --- | --- | --- | --- | --- | --- | --- | --- |
| Stroke | 4354 | 0 | 0.217 | 1215 | 0.217 | 120 | 0.261 |
| IscStroke | 3308 | 0 | 0.312 | 1266 | 0.312 | 202 | 0.333 |
| SAHStroke | 665 | 0 | NA | 722 | NA | 69 | NA |
| ICHStroke | 581 | 2 | NA | 1059 | NA | 146 | NA |

### 2.2. Evaluation of QTPhenProxy Model Performance

To evaluate the performance of the QTPhenProxy model, we determined its ability to recover cases when the defining ICD10 codes are removed. We trained 50% of the cases and an equal number of controls on the clinical features described above using the EN and RF classifiers. We chose these classifiers because of their overall high performance and their probability assignment distributions were continuous (Figure 1, Supplementary Figure A.1). We removed the ICD10 codes used to define the cases and controls in our feature set. We then tested the model on all other subjects in the UK Biobank, which included known cases and self-reported cases that did not have an ICD10 code for the disease. We then evaluated the recovery of 1) cross-validation and 2) the holdout test set through Precision at top 50, 100, 500, and number of cases, area under the receiver operating curve, area under the precision recall curve, and maximum F1 score.

**Figure 1:**
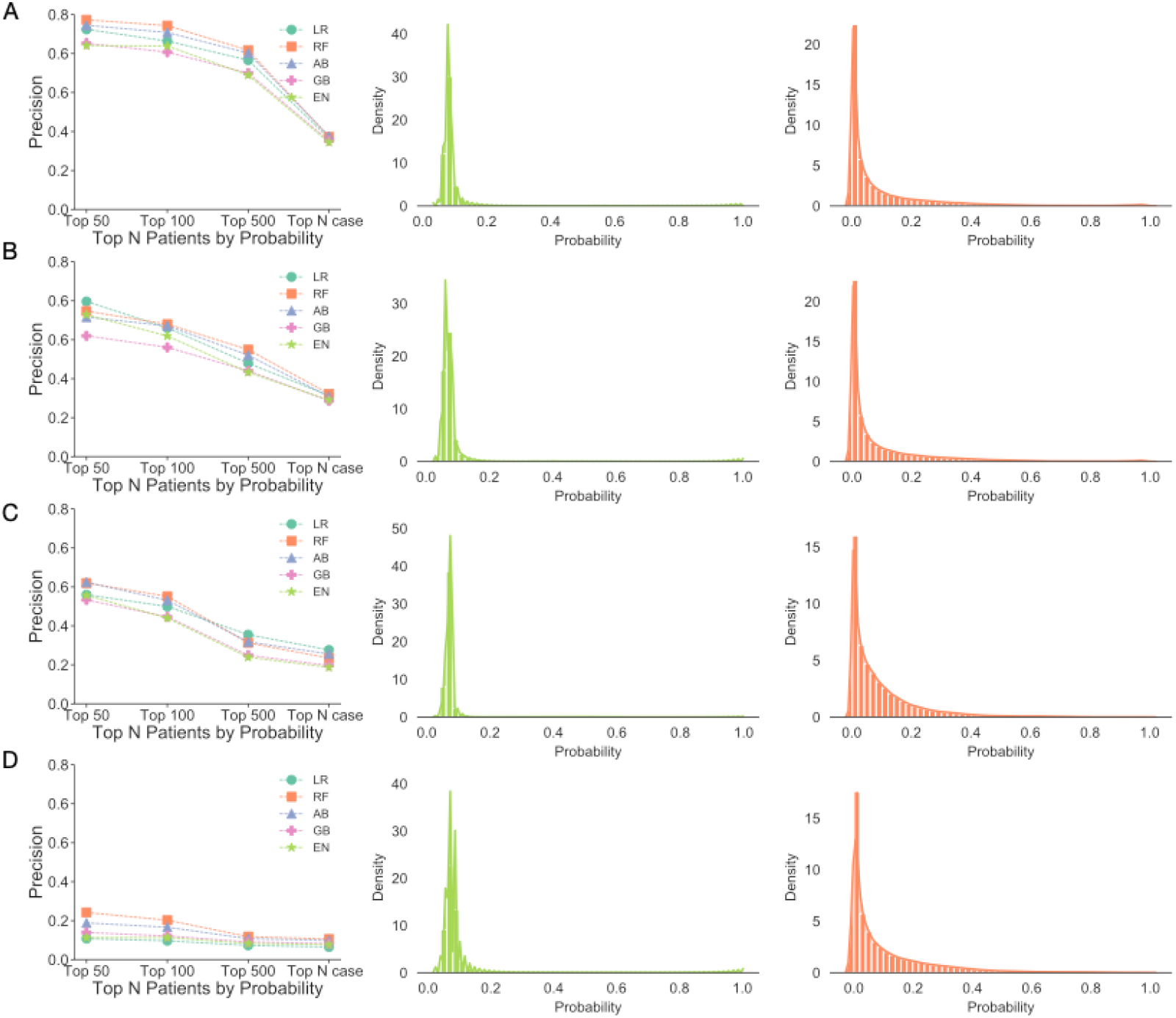
Precision at top 50, 100, 500, and N cases probabilities assigned by machine learning algorithms on hold out test set and probability distributions. A. All Stroke, B. Ischemic Stroke, C. Sub-arachnoid Hemorrhage, D. Intracerebral Hemorrhage. Left panel: Precision of each algorithm Circles represent precision for the Logistic Regression model with L1 penalty (LR), squares: Random Forest model (RF), triangles: Adaboost model (AB), cross: Gradient boosting model (GB), and star Logistic Regression model with Elastic Net penalty (EN). Middle Panel: EN probability distributions, Right panel: RF Probability distributions

### 2.3. Genotyping and Imputation

UK Biobank subjects that were of White British descent, in the UK Biobank PCA calculations and therefore without 3rd degree and above relatedness, and without aneuploidy were used in this study, totalling 337147 subjects (181,032 females and 156,115 males) (UK Biobank; Bycroft et al., 2018). Of the nearly 500,000 participants, approximately 50,000 subjects were genotyped on the UK BiLEVE Array by Affymetrix while the rest were genotyped using the Applied Biosystems UK Biobank Axiom Array, with over 800,000 markers. The arrays share 95% marker coverage. Initially, we ran a QC extracting markers with an MAF>0.5%, INFO score > 0.3, and hardy-weinberg equilibrium test mid-p value > 10^−10^ using all subjects, which we will refer to as QC1. QC1 was run for GWAS of all 18 diseases. We then re-ran the QC, which we will refer to as QC2, more stringently by extracting markers with an MAF>1%, INFO score > 0.8, and hardy-weinberg equilibrium test mid-p value > 10^−5^ using Plink2 (Chang et al., 2015). UKBB version 3 Imputation combined the Haplotype Research Consortium with the UK10K haplotype resource using the software IMPUTE4 (UK Biobank).

### 2.4. Genome-wide Association Analysis

The binary trait and QTPhenProxy probabilities were compared by running two separate association analyses. For both analyses, covariates included age at 2018, sex, first 10 principal components, and the genotyping array the sample was carried out on. In QC1, the original PCs determined by the UKBiobank QC were used. In QC2, we calculated the PCs using the method described in 2.11. QC2 GWAS was only run for stroke, ischemic stroke, subarachnoid hemorhage, and intracerebral hemorrhage GWAS. For the binary trait GWAS, a logistic regression was run adjusted with the aformentioned covariates. For comparison, the QTPhenProxy probabilities were quantile normalized and run under a linear regression adjusting for the same covariates. We also permuted the probabilities within the phenotyping files and ran additional GWAS 10 times to ensure the signal was correlated with the phenotype.

### 2.5. Mapping variants to known disease variant marker sets and mapping marker sets to disease systems

The EBI-GWAS catalogue has a database of all published GWAS (Buniello et al., 2019). We extracted over 2,000 disease marker sets conducted on populations with European ancestry. MedDRA is a standardized medical vocabulary developed by International Council for Harmonisation of Technical Requirements for Pharmaceuticals for Human Use (ICH) (Brown, 1999). All terms in the vocabulary can be mapped to its highest system level, which includes 27 different organ systems and other general and lab studies such as social circumstance and investigations. Using the NCBO annotator, we mapped the names of the EBI-GWAS disease marker sets to the MedDRA System Organ Classes level (Whetzel et al., 2011).

### 2.6. Assessing the specificity of the QTPhenProxy-derived variants

To assess the disease specificity of the genome-wide significant variants, we first calculated the proportion of genome-wide significant variants in each of the EBI-GWAS disease marker sets. We then aggregated the marker sets together by System Organ Class to evaluate the systems enriched for genome-wide significant variants. We ordered the marker sets in each class by proportion of genome-wide significant variants and divided by the number of marker sets in each class. We also compared the proportion of variants of varying significance of the marker sets related to each disease with 1) the other marker sets related to the same System Organ Class as the disease and 2) all other marker sets. We stratified each comparison by significance value, between 0.05 and 5E-08.

### 2.7. Evaluation of recovery of known variants

For each disease, we gathered EBI-GWAS marker sets that contained the disease in its name. These represent known variants of each disease. We then extracted the p-value from either the binary trait logistic regression or QTPhenProxy linear regression. We then ran a t-test comparing the negative log base 10 p-values of the binary trait with QTPhenProxy GWAS. We also ran a t-test comparing the difference between the binary and QTPhenProxy log base 10 p values for the known ischemic stroke variants and an equivalent number of random variants.

### 2.8. Refinement of discovered variants by QTPhenProxy using conditional analysis

At each LD-independent locus, the SNP with lowest p-value may not be the variant that causes the most phenotypic variation within the area (Yang et al., 2012). Therefore, we applied GCTA-COJO, a conditional analysis that takes into account lead SNPs and the LD structure of a sample of the population, to our genome-wide association results (Yang et al., 2012). We randomly sampled 10,000 subjects from the UKBiobank for the linkage disequilibrium calculation (Wells et al., 2019). From the GCTA-COJO results, we then mapped each locus to its nearest gene using dbSNP and the UCSC Genome Browser accessed at http://genome.ucsc.edu/ (National Center for Biotechnology Information, National Library of Medicine; Kent et al., 2002). For intergenic loci, we chose the 1-2 nearest genes that were at most 10,000 KB away.

### 2.9. Correlation of QTPhenProxy GWAS beta coefficients to Binary trait GWAS Odds Ratio

We calculated the Pearson correlation between the beta-coefficients of the QTPhenProxy GWAS and log of the odds ratios of the Binary trait GWAS. In order to account for noise, we calculated the correlation with variants with different levels of significance in the QTPhenProxy models. P-value cutoffs included 1, 0.05,0.0005,5e-06,and 5e-08.

### 2.10. Simulation of Conversion of QTPhenProxy trait to Binary trait and Conversion of beta coefficients to odds ratios

Based on the scale difference between binary and quantitative traits, the odds ratios are not directly comparable. We decided to simulate a method to convert beta coefficients from QTPhenProxy to Odds ratios. We did this by simulating a quantitative trait, converting it to a binary trait, and testing the correlation of a simulated marker variant's significance between the two methods. Using SOLAR, a software package for estimating heritability using identity by descent calculations, we simulated a quantitative trait with one quantitative trait locus with two alleles and a nearby marker locus with two alleles (Almasy and Blangero, 1998). We first removed all related individuals with a resulting cohort of 4,195 subjects. For the simulation, we varied the frequency for the causal minor allele and a marker minor allele from 0.05-0.45 in increments of 0.010, the mean quantitative trait value for the heterozygous genotype from 5-45 in increments of 10 and the homozygous genotypes' *mean* ± 50, the standard deviation of the quantitative trait from 5-20 in increments of 4, and the recombinant fraction from 0.01-0.10 in increments of 0.02. After simulating the quantitative trait distribution, we then normalized the trait to several distributions: standard normal, normal distribution with mean 0 and standard deviation 10, and mean 50 and standard deviation 10. We compared distributions because we quantile normalized our QTPhenProxy trait values before running the genome-wide associations studies. We then converted each simulation to a binary trait using liability thresholding (Risch, 1990). Liability thresholding was implemented as follows: We determined a quantitative trait value as a threshold based off the prevalence of the simulated trait, which we varied from 2.5-20% in increments of 5%. Any subjects above this threshold is labeled a case, and the rest controls in the binary trait phenotype. We then ran linear or logistic regressions using the python package statsmodels between the simulated quantitative trait or the binary trait and the subjects’genotypes for the marker and causal loci (Seabold and Perktold, 2010). We developed a conversion formula for the beta coefficients to odds ratios by linearly regressing the correlation between the simulated effect sizes. We then converted the beta-coefficients to odds ratios of the UK Biobank GWAS results by multiplying the beta coefficient by the average slope and intercept and then taking the exponential of the result.

### 2.11. PCA

In order to confirm the cases were well distributed within the data and to determine the number of principal components to use as covariates, we conducted PCA. For each of the four diseases, we first pruned the variants used for PCA by running a sliding window of size 100 kbps, 5 variant step size, and *r*^2^ threshold of 0.1. We then combined the chromosomes, extracting only the pruned variants, using plink2 and cat-bgen software (Band and Marchini, 2018). We then ran pca with plink2 (Chang et al., 2015) and evaluated the PCs to be used as covariates using a skree plot. Within the main PCs, we plotted them against each other, highlighting the distribution of the cases.

### 2.12. LD Score Regression and evaluation of genomic inflation

We determined the lambda genomic correction and LD score regression coefficient using the software LDSC (Bulik-Sullivan et al., 2015). We used the disease GWAS summary statistics and European LD scores pre-computed from 1000 genomes by the Alkes group (Bulik-Sullivan et al., 2015). QQ plots were plotted using qqman (Turner, 2014). To determine the relationship between genomic inflation and minor allele frequency, we binned all variants in the stroke gwas by minor allele frequency into 30 bins. We then calculated the genomic inflation of the p-values of the variants in each bin.

### 2.13. Genetic Correlation of QTPhenProxy with MEGASTROKE and Coronary Artery Disease GWAS

We measured the genetic correlation of the QTPhenProxy EN stroke and stroke subtype models using both QC1 and QC2 quality control and the results from the binary classification of stroke with a large stroke study as well as other stroke-related risk factors using LDSC software (Bulik-Sullivan et al., 2015). We gathered summary statistics from the MEGASTROKE meta-analysis of GWA data on stroke and stroke subtypes, including Acute Ischemic Stroke (AIS) (Malik et al., 2018a). We compared our results to these summary statistics and measured its genetic correlation with the same risk factor GWAS compared in MEGASTROKE (Malik et al., 2018a). Specifically, we measured the genetic correlation with the following risk factors: Coronary Artery Disease (CAD) (Nikpay et al., 2015), T2D (Xue et al., 2018), Atrial Fibrillation (AF) (Nielsen et al., 2018), White matter hyperintensities (WMH) (Traylor et al., 2019), and Low-density lipoprotein (LDL) (Willer et al., 2013). Further information about the resources of these risk factors can be found in Supplementary Information. Finally, we used LD Hub (Zheng et al., 2017), a database server by the Broad Institute that calculates genetic correlations with hundreds of available traits. To measure the genetic correlation between the QTPhenProxy EN stroke model and a wide variety of trait GWAS, from stroke-related to many that are unrelated to the pathophysiology of stroke, we ran LD Hub to calculate the genetic correlation between the QTPhenProxy Stroke QC1 GWAS and 597 trait GWAS run by the Neale lab in the UK Biobank cohort (Neale, 2018).

## 3. Results

### 3.1. QTPhenProxy has high model performance

We trained models with 5 different classifier types and all stroke, ischemic stroke, intracerebral hemorrhage, and subarachnoid hemorrhage as defined by the UK Biobank Algorithmically Defined Outcomes rubric (Schnier et al., 2017). Random forest models overall showed the best area under the receiver operating curve and adaboost models gave the best area under the precision recall curve on the hold-out test set (Supplementary Tables B.1 and B.3). Overall stroke, followed by ischemic stroke showed high precision at top 50 patients ordered by probability (Figure 1, Table 1, Supplementary Table B.2). We chose the probabilities from the EN and RF models to run the genome-wide association analyses because the distribution of their probabilities was continuous and included values from 0-1 (Figure 1, Supplementary Figure A.1).

### 3.2. Novel and known variants are recovered by QTPhenProxy in higher numbers traditional binary method using the QC1 markers and principal components

The binary trait logistic regression genome-wide association study for stroke, ischemic stroke, and subarachnoid hemorrhage recovered no variants with genome-wide significance. Binary trait logistic regression genome-wide association study for intracerebral hemorrhage recovered 2 variants with genome-wide significance. Quantitative trait linear regression using QTPhenProxy probabilities for stroke recovered 120 genome-wide significant SNPs with 16 LD-independent loci and 1215 genome-wide significant SNPs with 39 LD-independent loci, for ischemic stroke recovered 202 genome-wide significant SNPs with 15 LD-independent loci, and 1266 genome-wide significant SNPs with 46 LD-independent loci, for subarachnoid hemorrhage recovered 69 genome-wide significant SNPs with 4 LD-independent loci, and 722 genome-wide significant SNPs with 40 LD-independent loci, and for intracerebral hemorrhage recovered 146 genome-wide significant SNPs with 8 LD-independent loci, and 1059 genome-wide significant SNPs with 54 LD-independent loci using the EN and RF classifiers, respectively (Table 1). We show the comparison of genome-wide significant variants between the QTPhenProxy EN model and Binary trait GWAS for stroke in a Hudson plot (Figure 2) (Lucas). Out of known ischemic stroke variants, both models also recovered 3 known variants with genome-wide significance, and the EN model recovered 21/49 of the known variants, equivalent to a sensitivity of 0.428, while the traditional binary trait recovered 15/49, (sensitivity=0.306), and RF model recovered 14/49 (sensitivity=0.286) with nominal p-value of 0.05 (Table 1). For all stroke, sensitivity of known stroke variants was 0.333 using the EN model, 0.261 using the RF model, and 0.202 using the binary trait. Subarachnoid hemorrhage did not have a specific EBI-GWAS disease marker set, and the EBI-GWAS disease marker set for intracerebral hemorrhage only consisted of one variant. The difference in p-values between the binary and QTPhenProxy for known ischemic stroke variants was significantly increased compared to the same number of random variants using the EN classifier (two sample t test, t=2.43, p=0.0184 for EN classifier and t=1.74, p=0.0876 for RF classifier). In addition, QTPhenProxy with EN classifier showed a significant decrease in p-value of all known ischemic stroke variants compared to traditional binary method (two sample t-test, t=-2.1 p=0.0367) while QTPhenProxy with RF classifier did not show a significant decrease (t=-1.48,p=0.144). The difference in p-values between the binary and QTPhenProxy for known all stroke variants was significantly increased compared to the same number of random variants for the EN classifier (t=2.80, p=0.00638) but not significant for the RF classifier (t=0.737,p=0.463). In addition, QTPhenProxy with EN classifier showed a decrease in p-value of all known stroke variants compared to traditional binary method (two sample t-test, t=-1.87 p=0.0639) while QTPhenProxy with RF classifier did not show a significant decrease (t=-1.05,p=0.298).

**Figure 2:**
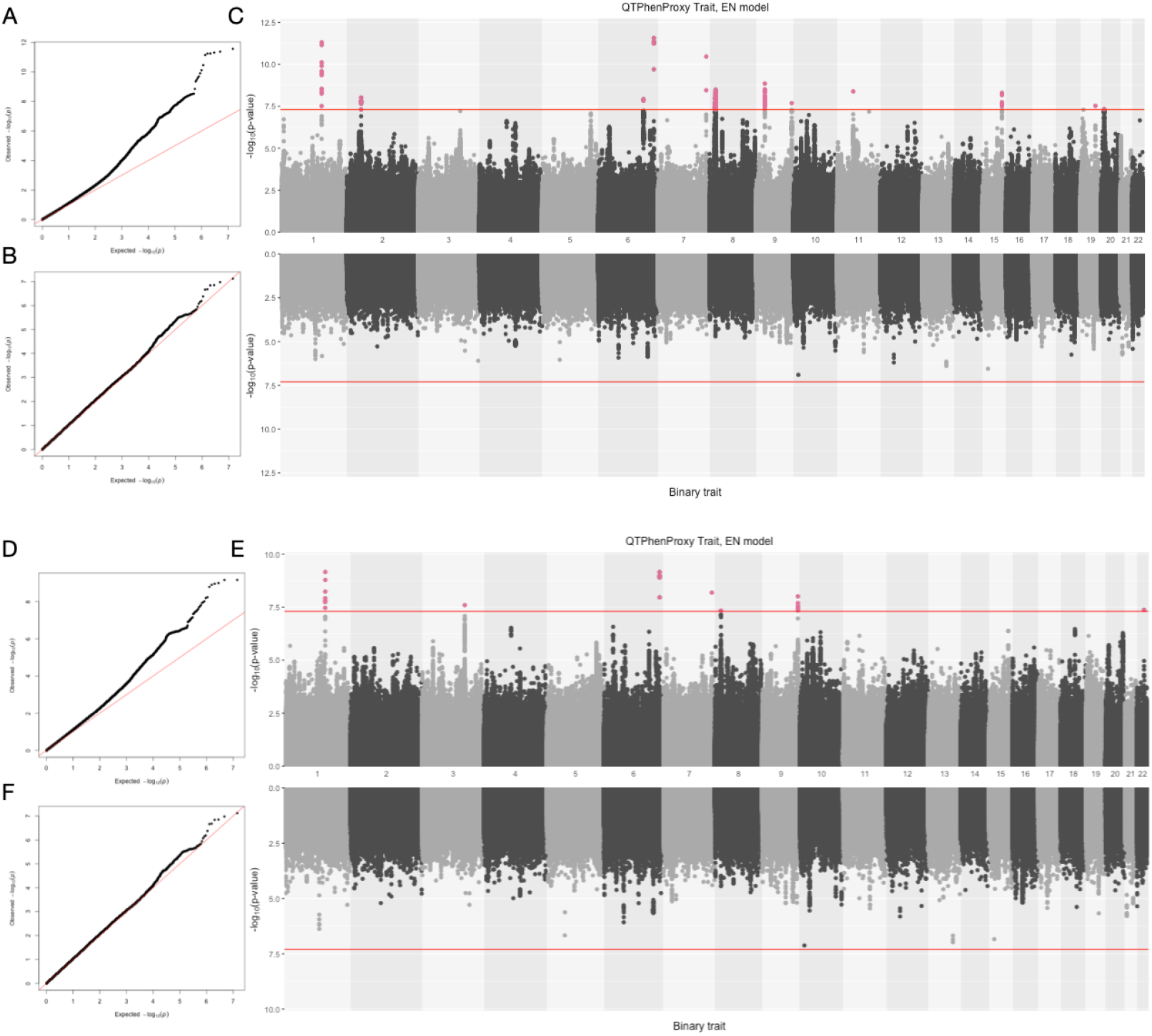
QQ Plots and Hudson plot of QTPhenProxy genome-wide association analysis, EN Model with Binary trait genome-wide association analysis for Stroke, (Top) using QC1 quality control and (Bottom) using QC2 quality control. A. and D. Q-Q Plot for QTPhenProxy, EN Model GWAS. B.and E. Q-Q Plot for Binary trait GWAS. C. and F. Top manhattan plot is for QTPhenProxy, bottom plot for binary trait.Variants with p-value < 5e-08 are highlighted in pink, and the dashed lines are at the same value.

### 3.3. Novel and known variants are recovered by QTPhenProxy in higher numbers traditional binary method using the QC2 markers and principal components

The binary trait logistic regression genome-wide association study for stroke, ischemic stroke, and subarachnoid hemorrhage recovered no variants with genome-wide significance. Binary trait logistic regression genome-wide association study for intracerebral hemorrhage recovered 2 variants with genome-wide significance. Quantitative trait linear regression using QTPhenProxy probabilities for stroke recovered 7 LD-independent loci, 3 LD-independent loci, for ischemic stroke, 3 LD-independent loci for subarachnoid hemorrhage, and 3 LD-independent loci for intracerebral hemorrhage using the EN classifier. We show the comparison of genome-wide significant variants between the EN and Binary in a Hudson plot (Figure 2) (Lucas). Out of known ischemic stroke variants, both models also recovered 3 known variants with genome-wide significance, and the EN model recovered 16/49 of the known variants, equivalent to a sensitivity of 0.326, while the traditional binary trait recovered 15/49, (sensitivity=0.306), and RF model recovered 15/49 (sensitivity=0.306) with nominal p-value of 0.05 (Figure 3). For all stroke, sensitivity of known stroke variants was 0.265 using the EN model, 0.220 using the RF model, and 0.220 using the binary trait (Supplementary Figure A.5). The t-tests referred to in 3.2 did not show significance.

**Figure 3:**
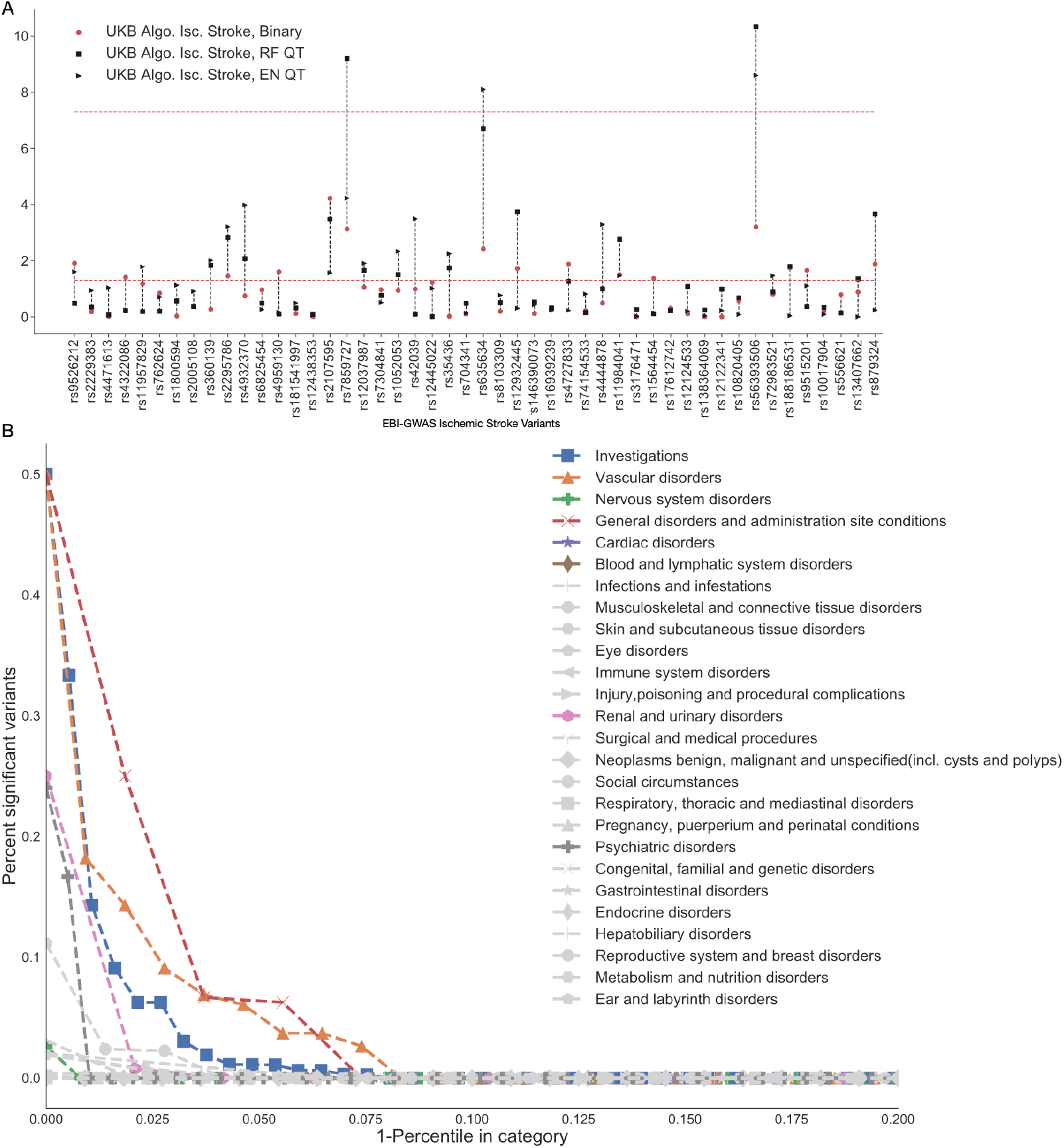
QTPhenProxy recovers known ischemic stroke variants and variants specific to related organ systems. Results from QTPhenProxy GWAS with QC2 quality control. (A) Horizontal axis shows Ischemic Stroke variants catalogued in EBI-GWAS that have shown genomewide significance in previous studies. Black markers represent p-values of variants recovered by QTPhenProxy models (square=RF,Triangle=EN). (B) X-axis is the top percentile of marker sets in each category, Y-axis is the proportion of variants in marker sets that overlap with QTPhenProxy genome-wide significant variants. Each shape corresponds to a disease category. In color are top disease categories: Square=Investigations, Triangle=Vascular disorders, Cross=Nervous system disorders, X=General disorders and administration site conditions, Star=cardiac disorders, and Diamond=Blood and lymphatic system disorders.

### 3.4. Conditional analysis identifies novel variants with genome-wide significance using the QC1 markers and principal components

Conditional analysis of the GWAS with QC1 quality control for the QTPhenProxy EN model for all stroke identified 13 candidate variants with genome-wide significance, all which are novel, though some of the nearest genes to these loci have been identified in previous studies through nearby loci. There were 12 loci identified for the stroke subtype ischemic stroke, 3 loci for subarachnoid hemorrhage, and 9 loci for intracerebral hemorrhage (Table 2). Several loci overlapped across stroke and some subtypes. Position 46705193 on chromosome 11, which is in an intronic section of *ARHGAP1* (National Center for Biotechnology Information, National Library of Medicine), showed genome-wide significance in stroke, ischemic stroke, and intracerebral hemorrhage. A missense mutation at locus rs6025 in the *F5* gene showed genome-wide significance across stroke, intracerebral hemorrhage, and locus rs1894692, which is near the *F5* gene (National Center for Biotechnology Information, National Library of Medicine), showed genome wide significance in ischemic stroke. Additional loci that showed genome-wide significance in both all stroke and ischemic stroke included those that are intronic to *LPA* and *NOS3* and nearby to *APOC1P1/APOC1* and *CDKN2A/CDKN2B* (National Center for Biotechnology Information, National Library of Medicine). Finally, all stroke and intracerebral hemorrhage shared additional loci that showed genome-wide significance after conditional analysis, which were nearby LOC105377992 and *RPS4X9* (National Center for Biotechnology Information, National Library of Medicine). All stroke genome-wide significant variants also included intronic loci in the *MTA3, SOX7, ABO, FURIN*, and *PLCB1* genes (National Center for Biotechnology Information, National Library of Medicine). Ischemic stroke genome-wide significant variants included intronic loci in theLPAL2 gene, intergenic loci in the *PITX2, LDLR*, and an insertion mutation in the *ABO* gene (National Center for Biotechnology Information, National Library of Medicine). Subarachnoid hemorrhage genome-wide significant variants included an intronic locus in the *ABO* gene. Finally, intracerebral hemorrhage genome-wide significant variants included a missense mutation loci in the *ABCG8* gene, and intronic loci in the *NOS3* and *XKR6* genes (National Center for Biotechnology Information, National Library of Medicine). The variants near or in LOC105377992, *NOS3, CDKN2A/CDKN2B, LPA, LDLR, ABCG8*, and *RPS4X9* were below nominal significance of 0.05 in the MEGASTROKE stroke and ischemic stroke GWAS of European ancestry (Table 3) (Malik et al., 2018a).

**Table 2:** Genome-wide significant variants discovered by QTPhenProxy, EN Model, using QC1 quality control. Cytogenic position was determined using Kent et al. (2002). *rsID*: variant id, *Chr*: cytogenic position, *Location*: location of variant relative to the gene, *R/R*: risk/reference, *RAF*: risk allele frequency in population, *Beta*: Beta coefficient OR: Odds Ratio, 95% Confidence Intervals

| Phenotype | rsID | Chr. | Gene | Location | R/R allele | RAF | Beta | Converted OR | P value |
| --- | --- | --- | --- | --- | --- | --- | --- | --- | --- |
| Stroke | rs6025 | 1q24.2 | *F5* | missense | T/C | 0.0231 | 0.0492 (0.0421-0.0564) | 1.09 (1.08-1.10) | **4.87e-12** |
| Stroke | rs8179838 | 2p21 | *MTA3* | intronic | T/C | 0.621 | 0.0126 (0.0104-0.0149) | 1.02 (1.02-1.03) | **9.63e-09** |
| Stroke | rs6923947 | 6q22.32 | *LOC105377992* | intergenic | A/G | 0.447 | 0.0123 (0.0101-0.0144) | 1.02 (1.02-1.02) | **1.19e-08** |
| Stroke | rs55730499 | 6q25.3 | *LPA* | intronic | T/C | 0.0817 | 0.0273 (0.0234-0.0312) | 1.05 (1.04-1.06) | **2.70e-12** |
| Stroke | rs3918226 | 7q36.1 | *NOS3* | intronic | T/C | 0.0814 | 0.0263 (0.0223-0.0302) | 1.05 (1.04-1.05) | **3.44e-11** |
| Stroke | rs6991641 | 8p23.1 | *SOX7* | intronic | G/C | 0.403 | 0.013 (0.0108-0.0152) | 1.02 (1.02-1.03) | **3.29e-09** |
| Stroke | rs1333049 | 9p21.3 | *CDKN2A-CDKN2B* | intergenic | C/G | 0.481 | 0.0129 (0.0108-0.0151) | 1.02 (1.02-1.03) | **1.41e-09** |
| Stroke | 9:136138765 | 9q34.3 | *ABO* | intronic | G/GC... | 0.184 | 0.0155 (0.0128-0.0183) | 1.03 (1.02-1.03) | **2.05e-08** |
| Stroke | 11:46705193 | 11p11.2 | *ARHGAP1* | intronic | C/CT | 0.28 | 0.0143 (0.0119-0.0167) | 1.02 (1.02-1.03) | **4.10e-09** |
| Stroke | rs59065675 | 15q26.1 | *FURIN* | intronic | C.../C | 0.47 | 0.0127 (0.0105-0.0149) | 1.02 (1.02-1.03) | **5.00e-09** |
| Stroke | rs814573 | 19q13.32 | *APOC1-APOC1P1* | intergenic | T/A | 0.188 | 0.0156 (0.0128-0.0184) | 1.03 (1.02-1.03) | **2.97e-08** |
| Stroke | 20:862958 | 20p12.3 | *PLCB1* | intronic | TA/T | 0.76 | 0.0138 (0.0113-0.0164) | 1.02 (1.02-1.03) | **4.73e-08** |
| IscStroke | rs1894692 | 1q24.2 | *F5-SLC19A2* | intergenic | G/A | 0.0218 | 0.0546 (0.0478-0.0614) | 1.1 (1.09-1.11) | **8.22e-16** |
| IscStroke | rs369787256 | 4q25 | *PITX2-MIR297* | intergenic | C/T | 0.0905 | 0.0223 (0.0188-0.0257) | 1.04 (1.03-1.05) | **1.17e-10** |
| IscStroke | rs55730499 | 6q25.3 | *LPA* | intronic | T/C | 0.0817 | 0.0272 (0.0236-0.0307) | 1.05 (1.04-1.05) | **2.49e-14** |
| IscStroke | rs117733303 | 6q25.3 | *LPAL2* | intronic | G/A | 0.0187 | 0.0428 (0.0356-0.05) | 1.08 (1.06-1.09) | **2.69e-09** |
| IscStroke | rs3918226 | 7q36.1 | *NOS3* | intronic | T/C | 0.0814 | 0.0209 (0.0173-0.0246) | 1.04 (1.03-1.04) | **6.81e-09** |
| IscStroke | rs2205258 | 8q23.3 | *LOC107986968* | intronic | C/T | 0.83 | 0.0151 (0.0125-0.0177) | 1.03 (1.02-1.03) | **6.76e-09** |
| IscStroke | rs8176719 | 9q34.2 | *ABO* | insertion | TC/T | 0.34 | 0.0145 (0.0124-0.0165) | 1.02 (1.02-1.03) | **2.17e-12** |
| IscStroke | rs1333049 | 9p21.3 | *CDKN2A-CDKN2B* | intergenic | C/G | 0.481 | 0.0132 (0.0112-0.0151) | 1.02 (1.02-1.03) | **1.31e-11** |
| IscStroke | 11:46705193 | 11p11.2 | *ARHGAP1* | intronic | C/CT | 0.28 | 0.0122 (0.01-0.0145) | 1.02 (1.02-1.02) | **3.22e-08** |
| IscStroke | rs814573 | 19q13.32 | *APOC1-APOC1P1* | intergenic | T/A | 0.188 | 0.0172 (0.0146-0.0198) | 1.03 (1.02-1.03) | **1.85e-11** |
| IscStroke | rs118068660 | 19p13.2 | *LDLR* | intergenic | C/T | 0.901 | 0.0193 (0.0159-0.0227) | 1.03 (1.03-1.04) | **1.43e-08** |
| IscStroke | rs12151925 | 20p12.2 | *LOC101929413-FAT1P1* | intergenic | C/T | 0.222 | 0.0129 (0.0105-0.0152) | 1.02 (1.02-1.03) | **4.26e-08** |
| SAHStroke | rs6025 | 1q24.2 | *F5* | missense | T/C | 0.0231 | 0.0673 (0.0593-0.0753) | 1.12 (1.11-1.14) | **4.61e-17** |
| SAHStroke | rs36058710 | 9q34.2 | *ABO* | intronic | CT/C | 0.287 | 0.0166 (0.0139-0.0194) | 1.03 (1.02-1.03) | **1.03e-09** |
| SAHStroke | rs34850248 | 12q12 | *LINC02400* | intergenic | A/AC | 0.0714 | 0.0268 (0.0221-0.0314) | 1.05 (1.04-1.06) | **9.95e-09** |
| ICHStroke | rs6025 | 1q24.2 | *F5* | missense | T/C | 0.0231 | 0.0480 (0.0411-0.0549) | 1.09 (1.07-1.1) | **2.85e-12** |
| ICHStroke | rs11887534 | 2p21 | *ABCG8* | missense | G/C | 0.935 | 0.0271 (0.0229-0.0313) | 1.05 (1.04-1.06) | **7.90e-11** |
| ICHStroke | rs372302634 | 3q22.3 | *ESYT3-MRAS* | intergenic | T/T... | 0.457 | 0.0121 (0.01-0.0142) | 1.02 (1.02-1.02) | **8.51e-09** |
| ICHStroke | rs6923947 | 6q22.32 | *LOC105377992* | intergenic | A/G | 0.447 | 0.0127 (0.0106-0.0148) | 1.02 (1.02-1.03) | **9.5e-10** |
| ICHStroke | rs2077111 | 6q22.33 | *RPS4XP9* | intergenic | G/A | 0.447 | 0.012 (0.00995-0.0141) | 1.02 (1.02-1.02) | **8.89e-09** |
| ICHStroke | rs1808593 | 7q36.1 | *NOS3* | intronic | G/T | 0.25 | 0.0131 (0.0107-0.0155) | 1.02 (1.02-1.03) | **4.14e-08** |
| ICHStroke | rs7013277 | 8p23.1 | *XKR6* | intronic | C/A | 0.396 | 0.0116 (0.00947-0.0137) | 1.02 (1.02-1.02) | **4.6e-08** |
| ICHStroke | 11:46705193 | 11p11.2 | *ARHGAP1* | intronic | C/CT | 0.28 | 0.0133 (0.011-0.0156) | 1.02 (1.02-1.03) | **1.35e-08** |

**Table 3:** Genome-wide significant variants discovered by QTPhenProxy, EN Model, using QC2 quality control. Also includes QC2 p-value for genome-wide significant variants from QC1 GWAS and p-values for variants from the MEGASTROKE stroke or ischemic stroke european GWAS Malik et al. (2018a). Cytogenic position was determined using Kent et al. (2002). *rsID*: variant id, *Chr*: cytogenic position

| Phenotype | rsID | Chr. | Gene | P value | MEGASTROKE P value |
| --- | --- | --- | --- | --- | --- |
| Stroke | rs1894692 | 1q24.2 | *F5-SLC19A2* | **6.87E-10** | 0.435 |
| Stroke | rs8179838 | 2p21 | *MTA3* | 8.36E-06 | 0.515 |
| Stroke | rs769407520 | 3q22.3 | *ATP5MC1P3* | **2.50E-08** | not found |
| Stroke | rs6923947 | 6q22.32 | *LOC105377992* | 4.58E-07 | 0.04332 |
| Stroke | rs55730499 | 6q25.3 | *LPA* | **6.85E-10** | 0.05915 |
| Stroke | rs3918226 | 7q36.1 | *NOS3* | **6.45E-09** | 0.00134 |
| Stroke | 8:10206921 | 8p23.1 | *MSRA* | **4.62E-08** | not found |
| Stroke | rs1333049 | 9p21.3 | *CDKN2A-CDKN2B* | 5.02E-06 | 1.85E-07 |
| Stroke | 9:136138765 | 9q34.3 | *ABO* | **9.73E-09** | not found |
| Stroke | 11:46705193 | 11p11.2 | *ARHGAP1* | 7.06E-07 | not found |
| Stroke | rs59065675 | 15q26.1 | *FURIN* | 4.06E-07 | not found |
| Stroke | rs814573 | 19q13.32 | *APOC1-APOC1P1* | 2.52E-05 | 0.412 |
| Stroke | 20:862958 | 20p12.3 | *PLCB1* | 1.95E-06 | not found |
| Stroke | rs2269608 | 22q13.1 | *KCNJ4* | **4.23E-08** | 0.212 |
| IscStroke | rs1894692 | 1q24.2 | *F5-SLC19A2* | **1.90E-11** | 0.229 |
| IscStroke | rs369787256 | 4q25 | *PITX2-MIR297* | 3.69E-06 | not found |
| IscStroke | rs55730499 | 6q25.3 | *LPA* | **1.26E-09** | 0.0316 |
| IscStroke | rs117733303 | 6q25.3 | *LPAL2* | 0.000119 | 0.422 |
| IscStroke | rs3918226 | 7q36.1 | *NOS3* | 3.59E-06 | 0.00118 |
| IscStroke | rs2205258 | 8q23.3 | *LOC107986968* | 7.16E-06 | 0.415 |
| IscStroke | 9:1361387 | 9q34.2 | *ABO* | **4.99E-10** | not found |
| IscStroke | rs1333049 | 9p21.3 | *CDKN2A-CDKN2B* | 3.01E-06 | 1.09E-06 |
| IscStroke | 11:46705193 | 11p11.2 | *ARHGAP1* | 3.85E-05 | not found |
| IscStroke | rs814573 | 19q13.32 | *APOC1-APOC1P1* | 2.20E-06 | 0.716 |
| IscStroke | rs118068660 | 19p13.2 | *LDLR* | 1.28E-06 | 1.14E-05 |
| IscStroke | rs12151925 | 20p12.2 | *LOC101929413-FAT1P1* | 0.000107 | 0.104 |
| SAHStroke | rs6025 | 1q24.2 | *F5* | **6.34E-18** | 0.453 |
| SAHStroke | rs36058710 | 9q34.2 | *ABO* | **1.59E-10** | not found |
| SAHStroke | rs34850248 | 12q12 | *LINC02400* | **7.18E-09** | not found |
| ICHStroke | rs6025 | 1q24.2 | *F5* | **1.52E-10** | 0.453 |
| ICHStroke | rs11887534 | 2p21 | *ABCG8* | **4.29E-08** | 6.85E-04 |
| ICHStroke | rs372302634 | 3q22.3 | *ESYT3-MRAS* | **4.03E-08** | not found |
| ICHStroke | rs6923947 | 6q22.32 | *LOC105377992* | 1.47E-07 | 0.00433 |
| ICHStroke | rs2077111 | 6q22.33 | *RPS4XP9* | 1.96E-06 | 0.0372 |
| ICHStroke | rs1808593 | 7q36.1 | *NOS3* | 2.96E-06 | 0.565 |
| ICHStroke | rs7013277 | 8p23.1 | *XKR6* | 2.06E-05 | 0.156 |
| ICHStroke | 11:46705193 | 11p11.2 | *ARHGAP1* | 3.59E-06 | not found |

### 3.5. Conditional analysis identifies novel variants with genome-wide significance using the QC2 markers and principal components

Conditional analysis of the GWAS with QC2 quality control for the QTPhenProxy EN model for all stroke identified 7 candidate variants with genome-wide significance, all which are novel, though some of the nearest genes to these loci have been identified in previous studies through nearby loci. There were 3 loci identified for the stroke subtype ischemic stroke, 3 loci for subarachnoid hemorrhage, and 3 loci for intracerebral hemorrhage (Table 3). Almost all of the genome-wide significant variants overlapped with those found running the GWAS using QC1 quality control. New variants that were genome-wide significant for all stroke included a deletion variant that was intergenic to the gene *ATP5MC1P3*, an intronic variant of the *MRSA* gene, and a 3'UTR variant in *KCNJ4* (National Center for Biotechnology Information, National Library of Medicine). In addition, several genome-wide significant variants in the MEGASTROKE stroke and ischemic stroke GWAS were near the variants that were genome-wide significant in the QTPhenProxy EN GWAS. Almost all of those nearby MEGASTROKE genome-wide significant variants replicated to at least nominal significance of 0.05 in the QTPhenProxy EN GWAS (Table 4).

**Table 4:** QTPhenProxy EN Model GWAS using QC2 quality control P-value of variants that were genome-wide significant in MEGASTROKE Stroke and Ischemic Stroke GWAS.

| Phenotype | Gene | rsID | QTPhenProxy p-value |
| --- | --- | --- | --- |
| Stroke | *LPA* | rs56393506 | 1.09E-08 |
| Stroke | *NOS* | rs1799983 | not found |
| Stroke | *CDKN2B-CDKN2A* | rs7859727 | 6.94E-05 |
| Stroke | *ABO* | rs635634 | 1.07E-07 |
| Stroke | *FURIN* | rs4932370 | 2.55E-05 |
| Stroke | *COL4A1* | rs9521634 | 0.00375 |
| Stroke | *LOC105372798* | rs720470 | 0.88 |
| Ischemic stroke | *PITX2-MIR297* | rs13143308 | not found |
| Ischemic stroke | *LPA* | rs56393506 | 2.50E-09 |
| Ischemic stroke | *NOS3* | rs1799983 | not found |
| Ischemic stroke | *CDKN2B-CDKN2A* | rs7859727 | 5.95E-05 |
| Ischemic stroke | *ABO* | rs635634 | 7.93E-09 |
| Ischemic stroke | *LDLR* | rs8103309 | 0.171 |
| Subarachnoid Hemorrhage | *ABO* | rs635634 | 2.22E-06 |
| Intracerebral Hemorrhage | *NOS3* | rs1799983 | not found |

### 3.6. Effect sizes of QTPhenProxy and traditional binary trait analysis are correlated

We determined the correlation between the effect sizes of the binary trait GWAS and QTPhenProxy GWAS. Pearson correlation of the beta coefficients and log of the odds ratios increased when restricting to variants with small p-values (max *r*^2^=0.58 for GWAS with QC2 control, max *r*^2^=0.70 for GWAS with QC1 control) (Figure 4, Supplementary Figure A.6). Too few variants, such as with p-values > 5e-08 resulted in a decreased Pearson correlation.

**Figure 4:**
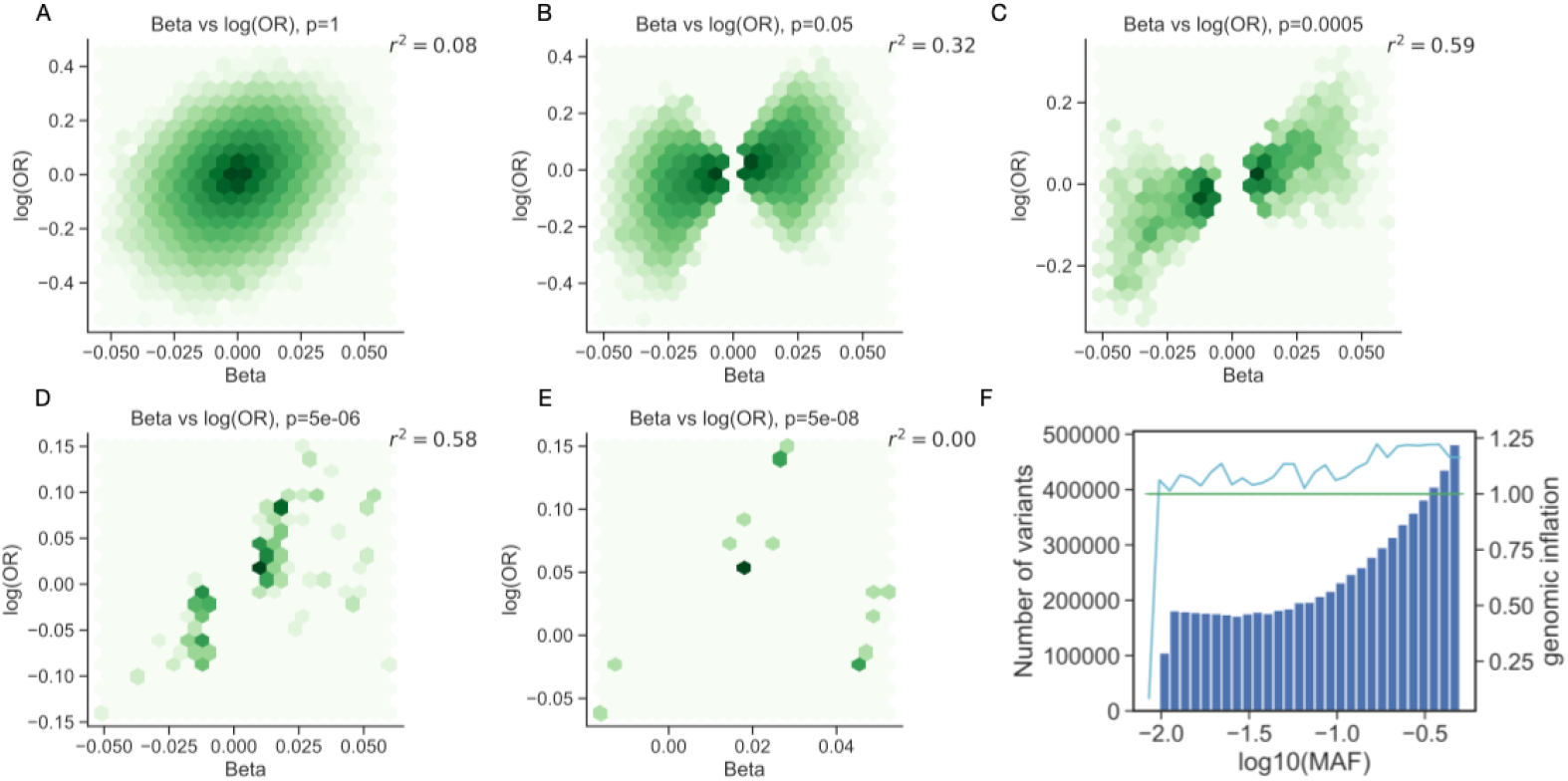
Correlation between beta and odds ratio effect sizes for variants at varying degrees of significance and Genomic Inflation within bins of variants with similar minor allele frequencies for QTPhenProxy EN Model for Stroke with QC2 quality control. (A-E) Correlation between QTPhenProxy, EN Model for Stroke GWAS and Binary trait GWAS effect size, QC2 quality control. X-axis is QTPhenProxy GWAS beta-coefficients, Y-axis is log base 10 of the Binary GWAS odds ratios. Pearson correlation is recorded in top right corner. From left to right, top to bottom, variants included decreases by restricting p-value.(F) Genomic Inflation within bins of variants with similar minor allele frequencies for QTPhenProxy EN Model for Stroke with QC2 quality control. Light blue line shows the genomic inflation of each bin of variants, green line shows 1.0 genomic inflation, and blue bars show numbers of SNPs (variants) in each bin, binned by *log10(MAF)*, or log(minor allele frequency.)

### 3.7. Genome-wide significant variants are enriched in related EBI-GWAS marker sets

From the over 2,000 EBI-GWAS disease variant marker sets mapped to organ systems, we calculated the proportion of markers in each set that was found to be at genome-wide significance by our QTPhenProxy model with QC2 markers and principal components. We found that the organ systems with markers sets with the highest proportion of genome-wide significant variants for stroke included vascular disorders, investigations, psychiatric disorders, and general disorders and administration site conditions (Figure A.5, Supplementary Figure 3). The enriched disease marker sets in the Investigations class included the lab values lipoprotein A levels, lipoprotein a levels adjusted for apolipoprotein A isoforms, blood protein levels, and white blood cell counts. The enriched disease marker sets in the General disorders and administration site conditions included aortic valve stenosis and allergy, while the Psychiatric disorders enriched marker sets were response to statins (LDL change) and venous thromboembolism. For ischemic stroke, the top enriched disease marker sets included those described for stroke and also activated partial thromboplastin time, coagulation factor levels, protein biomarkers, soluble levels of adhesion molecules, pancreatic cancer, and urinary metabolites.

### 3.8. LD score regression intercept and genomic inflation are similar across stroke and its subtypes

Using QC1 quality control, the genomic inflation for QTPhenProxy EN model for Stroke was 1.133, while using the QC2 quality control the genomic inflation for the same model was 1.134. For binary traits using QC2 quality control, genomic inflation for stroke was 1.027. Using the QC2 quality control model, the QTPhenProxy EN model genomic inflation for Ischemic stroke was 1.118, for subarachnoid hemorrhage was 1.122, and for intracerebral hemorrhage was 1.114. LD score regression intercept for QTPhenProxy EN model with QC2 quality control for stroke was 1.037 ± 0.0088, for ischemic stroke, 1.031 ± 0.0077, for subarachnoid hemorrhage, 1.012 ± 0.008, and for intracerebral hemorrhage, 1.051 ± 0.0081. We found that more common variants, or those with higher minor allele frequencies, had higher genomic inflation than rarer variants (Figure 4).

### 3.9. Genetic Correlation of QTPhenProxy Models' GWAS with MEGASTROKE GWAS and Risk Factor GWAS

We measured a genetic correlation of 0.60 (p-value 1.4E-19) between the QTPhenProxy EN Model for stroke, QC2 quality control, and the MEGASTROKE all stroke GWAS, and a genetic correlation of 0.49 (p-value 2.2E-12) between the QTPhenProxy EN Model for ischemic stroke, QC2 quality control, GWAS and the MEGASTROKE acute ischemic stroke GWAS (Malik et al., 2018a). We also found significant and large positive genetic correlations between QTPhenProxy EN Stroke and Ischemic Stroke Models and QTPhenProxy RF Stroke Model GWAS and coronary artery disease, type 2 diabetes mellitus, atrial fibrillation, and white matter hyperintensities (Supplementary Table B.4). We found significant and large negative correlations between subarachnoid hemorrhage and coronary artery disease, type 2 diabetes mellitus, atrial fibrillation and low density lipoprotein levels GWAS. Calculating genetic correlation calculations between the traditional binary GWAS for UK Biobank and MEGASTROKE GWAS and risk factors GWAS resulted in error due to low heritability (below 1E-03) estimated for the UK Biobank GWAS or invalid values encountered during genetic correlation calculation. After running genetic correlations on 1) the QTPhenProxy EN Model for stroke, QC1 quality control, GWAS and 597 trait GWAS run by the Neale lab on the UK Biobank data set, and 2) the QTPhenProxy EN Model for stroke, QC2 quality control, GWAS and 804 trait GWAS, including the same 597 UKBB traits, we found 117 and 118 traits, respectively, with bonferroni significant genetic correlations (see Supplementary Information). We found the top 50 significant genetic correlations for both models to be in traits associated with cardiovascular and kidney diseases, blood pressure and cardiovascular disease medications, anthropomorphic measurements such as waist circumference and BMI, physical activity, education, and family history (See Supplementary Information). We found the highest negative genetic correlations in traits that state no vascular or heart problems, no medications for cholesterol, blood pressure, or diabetes, and father's age at death. The insignificant and smallest magnitude of genetic correlations were with traits that included asthma, some cancer diagnoses, bone disorders and fractures, alcohol use, amyloidosis, asthma, type of transportation used, and asthma (See Supplementary Information).

### 3.10. Simulation of conversion of quantitative trait to binary trait shows correlation of effect sizes to empirical data

Based on the scale difference between binary and quantitative traits, the odds ratios are not directly comparable. We simulated the effect of a quantitative trait locus and nearby marker both on the original quantitative trait and the binary trait converted from the quantitative trait using liability thresholding. We found high correlation of effect sizes between the beta coefficients and log odds ratios of the quantitative trait variants with p-value < 0.005 and their respective binary trait variant effect sizes (Pearson correlation, *r*^2^=0.82). Lower p-values were not tested because of too few qualifying variants. After standard normalizing the quantitative trait, we also found the slope of the correlation between it and the log binary trait to be stable across the parameters, with a mean of 1.63 for each parameter except for marker allele frequency (1.35) and prevalence (1.77) (Supplementary Figure A.7).

## 4. Discussion

### 4.1. QTPhenProxy can identify patients with stroke using EHR data other than the disease diagnosis code

For all stroke and its subtype ischemic stroke, the machine learning models trained to assign QTPhenProxy probabilities performed well (greater than 90% AUROC, greater than 30% maximum F1 score, 74-97% precision at 50). The models trained on subarachnoid hemorrhage and intracerebral hemorrhage cases performed similarly with AUROC but not as well with maximum F1 score and precision at 50. This may be due to the number of cases the two subtypes have to train on, which was an order of magnitude smaller (600 cases versus 3300-4300 cases). Importantly, we only trained on half of the cases and tested on the other half, which equates to 300-2200 cases, depending on the disease definition. From this small training set, we were able to assign a probability of disease to all 500,000 subjects in the UK Biobank. This training set is a fraction of the over 40,000 cases required to power the most recent genome-wide association study for stroke (MEGASTROKE).

### 4.2. QTPhenProxy discovers many new variants and recovers known disease variants to genome-wide significance

Using QTPhenProxy resulted in 3-13 loci with genome-wide significance. In contrast, traditional binary trait GWAS using all disease cases resulted in the discovery of 0 genome-wide significant loci for stroke, ischemic stroke, and subarachnoid hemorrhage and 3 loci for intracerebral hemorrhage. In addition to new loci discovery, QTPhenProxy recovered known disease variants to a significance level of 0.05 with a sensitivity better than binary trait GWAS. In addition to recovering known variants, overall the effect size of the known variants in the binary trait GWAS was correlated with QTPhenProxy effect sizes for variants with low p-values. These results suggest that the QTPhenProxy method can recover relevant variants with fewer cases than traditional methods for stroke.

### 4.3. Simulation of quantitative trait and corresponding binary trait further supported the correlation of effect sizes between the two methods

Our simulations showed high correlation between quantitative trait beta coefficients and binary trait log odds ratio, which is similar to our empirical findings in the UK Biobank. We also show that the correlation slope is relatively stable across all the different simulation parameters, suggesting that there is a set correlation between quantitative traits and binary traits created from them. Further simulation will need to determine the effects of multiple loci on correlation between quantitative and corresponding binary trait.

### 4.4. Variants discovered for stroke are enriched in disease marker sets for vascular and neurological disease, and variants discovered for other diseases were enriched for disease and system specific markers

As a specificity measure for the variants discovered by QTPhenProxy, we found the EN model had the highest proportion of overlapping variants with EBI-GWAS marker sets related to vascular disorders and associated lab values. QTPhenProxy variants improves the power of detecting variants related to diseases that are co-morbid or risk factors for stroke.

### 4.5. QTPhenProxy has high genetic correlation with the MEGASTROKE GWAS and stroke-related risk factor GWAS

We found high genetic correlation (0.60-0.64) between the MEGASTROKE stroke and ischemic stroke GWAS and the QTPhenProxy models. The high genetic correlation between MEGASTROKE and QTPhenProxy suggests that both studies highlight shared genetic underpinnings, which we confirmed by finding large positive genetic correlations between the QTPhenProxy stroke and ischemic stroke models and risk factor GWAS that had large positive genetic correlations with MEGASTROKE (Malik et al., 2018a). We found large negative genetic correlations between the QTPhenProxy sub-arachnoid hemorrhage GWAS and the risk factors. We also were unable to find genetic correlations between the traditional binary GWAS of stroke, ischemic stroke, intracerebral hemorrhage, and sub-arachnoid hemorrhage and MEGASTROKE or the risk factors, most likely due to the small number of cases (581-4354) in each cohort. Finally, we found the least significant genetic correlations between the QTPhenProxy models and traits that are not stroke-related, such as bone disorders, cancer, and cataracts, while the most significant were cardiovascular-related. These findings suggest that our models do highlight genetic associations related to the pathophysiology of stroke. Although there is concern that this method may highlight genetic associations related to risk factors of stroke rather than stroke itself, we show that our models' GWAS are highly correlated with the MEGASTROKE GWAS and have similar correlations with risk factor GWAS highlighted by the study. These results suggest that QTPhenProxy models develop a phenotypic score ascertained for stroke and its subtypes.

### 4.6. Low LD score regression intercepts relative to genomic inflation suggests high polygenicity

Our genome-wide association studies using QTPhenProxy had moderate genomic inflation, slightly above 1.1, but low LD score regression intercepts near 1.0. The corresponding binary trait GWAS had genomic inflation below 1.05, suggesting minimal population stratification. Since the population for the binary trait GWAS was the same as those used for QTPhenProxy, this suggests that polygenicity, rather than population stratification, is the cause for genomic inflation. Polygenicity, or the contribution of small effects of many genes to a phenotype, may be the more likely cause (Bulik-Sullivan et al., 2015). In addition, Bulik-sullivan et. al. argue that genomic inflation can increase with sample size when there is polygenicity, and LD score regression intercept is more robust in distinguishing inflated p-values from polygenicity. As seen in (Howrigan et al., 2017), increased genomic inflation with common variants over rare variants suggests polygenicity. We show a similar upward trend in Figure 4. Another sign of true signal over inflated signal is the evidence of causal variants in linkage disequilibrium, as seen in a manhattan or hudson plot (Howrigan et al., 2017). Figure 2 shows genome-wide significant variants in linkage disequilibrium with each other. Finally, this study reports results from using two different quality controls in the genome-wide association analysis, QC1, and more stringent, QC2. Results using QC1 showed more variants with genome-wide significance and discovered more new variants for stroke than the QC2 GWAS. This may lead one to believe that more stringent quality control led to reduced inflation of p-values. However, the genomic inflation of the p-values from the QC1 GWAS was the same as the genomic inflation of the p-values from the QC2 GWAS (1.134 vs 1.133) with the QC2 GWAS having a slightly higher LD score regression intercept value (1.0288 vs 1.0369). This suggests that the p-values from the QC1 GWAS may not by overly inflated.

### 4.7. QTPhenProxy replicates known stroke variants and discovers variants within cardiovascular disease genes

Several of the genes discovered by the QTPhenProxy GWAS, using QC1 or QC2 quality control have been associated with stroke, including *NOS3*, *FURIN*, *PITX2, CDK2NB, LDLR*, and *ABO* (Malik et al., 2018a,b). *NOS3*, in particular was discovered through meta-analysis of the MEGASTROKE results with the UK Biobank (Malik et al., 2018b). We were able to replicate this association using only the QTPhenProxy method on the UK Biobank, and not traditional binary trait analysis. *NOS3* has been shown to be related to hypertension either through salt excretion regulation in the kidney (Gao Yang et al., 2018) or regulation of vascular relaxation in endothelial cells (Farah et al., 2018). Other discovered genes by QTPhenProxy are also associated with related cardiovascular diseases. *F5* gene codes for an essential coagulation factor, and mutations that can lead to increased thrombosis or hemorrhage, depending on the mutation (Asselta and Peyvandi, 2009; Kujovich, 2011; Hinds et al., 2016). *LPA* codes for a protein, lipoprotein-A, that can contribute to atherosclerosis (The Emerging Risk Factors Collaboration, 2009; Wang Long et al., 2016). Mutations in the *PLCB1* gene, which encodes for phospholipase synthesis, have been found in epilepsy and seizures (Schoonjans et al., 2016). *LDLR* gene codes for the low-density lipoprotein receptor, which is involved in cholesterol production (Brown and Goldstein, 1979), and the *APOC1/APOC1P1* genes code for parts of apoliprotein C1, which are involved in high density lipoprotein metabolism (Erqou et al., 2010). *ARHGAP1*, a gene coding for Rho GTPase activating protein 1 has been associated with cancer phenotypes and activation of hypoxic and inflammatory pathways (Fessler et al., 2004; Hashimoto et al., 2018; Zhang et al., 2019). *ABCG8* gene has been associated with combined gwas of lipids and inflammation, lipid levels, and gallstone disease (Ligthart et al., 2016). In addition, it is knownto code for a cholesterol transporter *EYST3* and *XKR6* gene mutations have been associated with coronary artery disease and ischemic stroke respectively in Asian populations, and *MRAS* gene mutations have been associated with coronary artery disease in two populations (Jiang et al., 2011; Zheng et al., 2019; Song et al., 2019; Erdmann et al., 2009). These results suggest that QTPhenProxy has replicated genome-wide significant mutations in genes known to be related with stroke and discovered of those associated with related risk factors such as coronary artery neurological diseases. Out of the 13 newly discovered variants with genome-wide significance, five are intronic or nearby genes that have been found in previous studies (MEGASTROKE), and five more are have been discovered in previous GWAS studies of related cardiovascular diseases. The variants within and near the *F5* gene had the highest genetic signal for stroke and all of its subtypes, even though it was not replicated in the MEGASTROKE GWAS. Out of all the new variants discovered by the QTPhenProxy EN model using the GWAS QC2 quality control, the rs11887534 variant, a missense mutation within the *ABCG8* gene, replicated to a p-value significance of 6.85E-04 in the MEGASTROKE stroke GWAS of over 40,000 European ancestry subjects. Using a tenth of the number of cases, QTPhenProxy discovered a variant within a new gene that replicated in MEGASTROKE, and discovered variants within known stroke genes.

### 4.8. Limitations

There are several limitations with this method. First, out of the five machine learning models used in QTPhenProxy, only two provided probabilities along a continuous scale. Adaboost, gradient boosting, and logistic regression using L1 penalty models, although with high performance, assigned probabilities in discrete bins. These distributions violate the the genome-wide association linear regression assumption of normal distribution of the quantitative trait. Although the probabilities produced by EN and RF models were not initially normally distributed, they were continuous, and could be adjusted with quantile normalization. In addition, the GWAS results from QTPhenProxy using the RF model resulted in many more hits than the EN model, and reduced sensitivity to known disease variants. Further study will be required to understand why one model gave more sensitive and specific results than the other, and whether there was p-value inflation in the random forest models. QTPhenProxy also may be particularly suited for stroke because the training model for phenotyping patients was optimized for stroke (Thangaraj et al., 2019). Since stroke is an acute event than can be identified with high accuracy in the electronic health record, this method may not translate as well to other diseases, such as chronic illnesses. A final limitation discussion point is whether QTPhenProxy models highlight genetic associations with risk factors for stroke rather than stroke itself. We have shown, from replicating known stroke variants to finding large positive and significant genetic correlations with MEGASTROKE and related traits, that our models improve the power of genetic associations for stroke, and not just a few specific risk factors.

### 4.9. Conclusions

We have developed a method, QTPhenProxy, that we have shown improves the power of genome-wide association studies in stroke and three of its subtypes: ischemic stroke, subarachnoid hemorrhage, and intracerebral hemorrhage with an order of magnitude fewer cases than required for traditional genome-wide association studies of the same diseases. Converting dichotomous traits to quantitative ones could result in improvement of power by incorporating electronic health record information for subjects who may have genetic susceptibility to stroke but may not have experience a stroke yet. Previous studies have shown that for diseases with low prevalence, there can be a reduction of power using logistic regression for binary trait GWAS compared to linear regression for quantitative trait GWAS (Zaitlen et al., 2012). Recently, (Abraham et al., 2019) showed that the inclusion of ischemic stroke risk factors' genome-wide significant SNPs in polygenic risk score improves prediction of ischemic stroke. This supports our idea that inclusion of risk factor information into the phenotype can help detect genetically susceptible subjects. We show that with as few as 2200 stroke subjects we can recover known variants of stroke and discover new variants that have been linked to cardiovascular and nervous system diseases. This method could be useful for studies with a small set of cases and without access to large meta-analyses. We also have suggested new variants that warrant further replication in other groups. QTPhenProxy shows the benefits of incorporating electronic health record data to convert traditional binary traits to quantitative in improving GWAS power.

## 5. Acknowledgements

All UK Biobank data analysis conducted in this work is available upon request. I would like to thank Dr. Krzysztof Kiryluk, Dr. Konrad Karczewski, Jie Yuan, Dr. Anna O. Basile for helpful discussions and guidance. Thank you also to Dr. Anna O. Basile, Nicholas Giangreco, and Jenna Kefeli for assistance in the UK Biobank application. This research has been conducted using the UK Biobank Resource under Application Number 41039. PT is funded by F30HL140946, and NT is funded by R35GM131905 and was funded by 5R01GM107145. MedDRAS^®^ trademark is registered by IFPMA on behalf of ICH. The MEGASTROKE project received funding from sources specified at http://www.megastroke.org/acknowledgments.html. We gratefully acknowledge all the studies and databases that made GWAS summary data available: ADIPOGen (Adiponectin genetics consortium), C4D (Coronary Artery Disease Genetics Consortium), CARDIoGRAM (Coronary ARtery DIsease Genome wide Replication and Meta-analysis), CKDGen (Chronic Kidney Disease Genetics consortium), dbGAP (database of Genotypes and Phenotypes), DIAGRAM (DIAbetes Genetics Replication And Meta-analysis), ENIGMA (Enhancing Neuro Imaging Genetics through Meta Analysis), EAGLE (EArly Genetics Lifecourse Epidemiology Eczema Consortium, excluding 23andMe), EGG (Early Growth Genetics Consortium), GABRIEL (A Multidisciplinary Study to Identify the Genetic and Environmental Causes of Asthma in the European Community), GCAN (Genetic Consortium for Anorexia Nervosa), GEFOS (GEnetic Factors for OSteoporosis Consortium), GIANT (Genetic Investigation of ANthropometric Traits), GIS (Genetics of Iron Status consortium), GLGC (Global Lipids Genetics Consortium), GPC (Genetics of Personality Consortium), GUGC (Global Urate and Gout consortium), HaemGen (haemotological and platelet traits genetics consortium), HRgene (Heart Rate consortium), IIBDGC (International Inflammatory Bowel Disease Genetics Consortium), ILCCO (International Lung Cancer Consortium), IMSGC (International Multiple Sclerosis Genetic Consortium), MAGIC (Meta-Analyses of Glucose and Insulin-related traits Consortium), MESA (Multi-Ethnic Study of Atherosclerosis), PGC (Psychiatric Genomics Consortium), Project MinE consortium, ReproGen (Reproductive Genetics Consortium), SSGAC (Social Science Genetics Association Consortium) and TAG (Tobacco and Genetics Consortium), TRICL (Transdisciplinary Research in Cancer of the Lung consortium), UK Biobank.

We gratefully acknowledge the contributions of Alkes Price (the systemic lupus erythematosus GWAS and primary biliary cirrhosis GWAS) and Johannes Kettunen (lipids metabolites GWAS).

## 6. Author Contributions

Conceptualization: P.T. and N.T., methodology: P.T and N.T., software: P.T. and N.T., formal analysis: P.T., U.G., and N.T., investigation: P.T. and N.T., resources: P.T. and N.T., data curation: P.T., U.G., and N.T., writing-original draft: P.T. and N.T., writing-review and editing: P.T., U.G., and N.T., visualization: P.T and N.T, Supervision: N.T., Funding Acquisition: P.T. and N.T.

## 7. Declaration of Interests

The authors declare no competing interests.

## Appendix A. Supplementary Figures

**Figure A.1:**
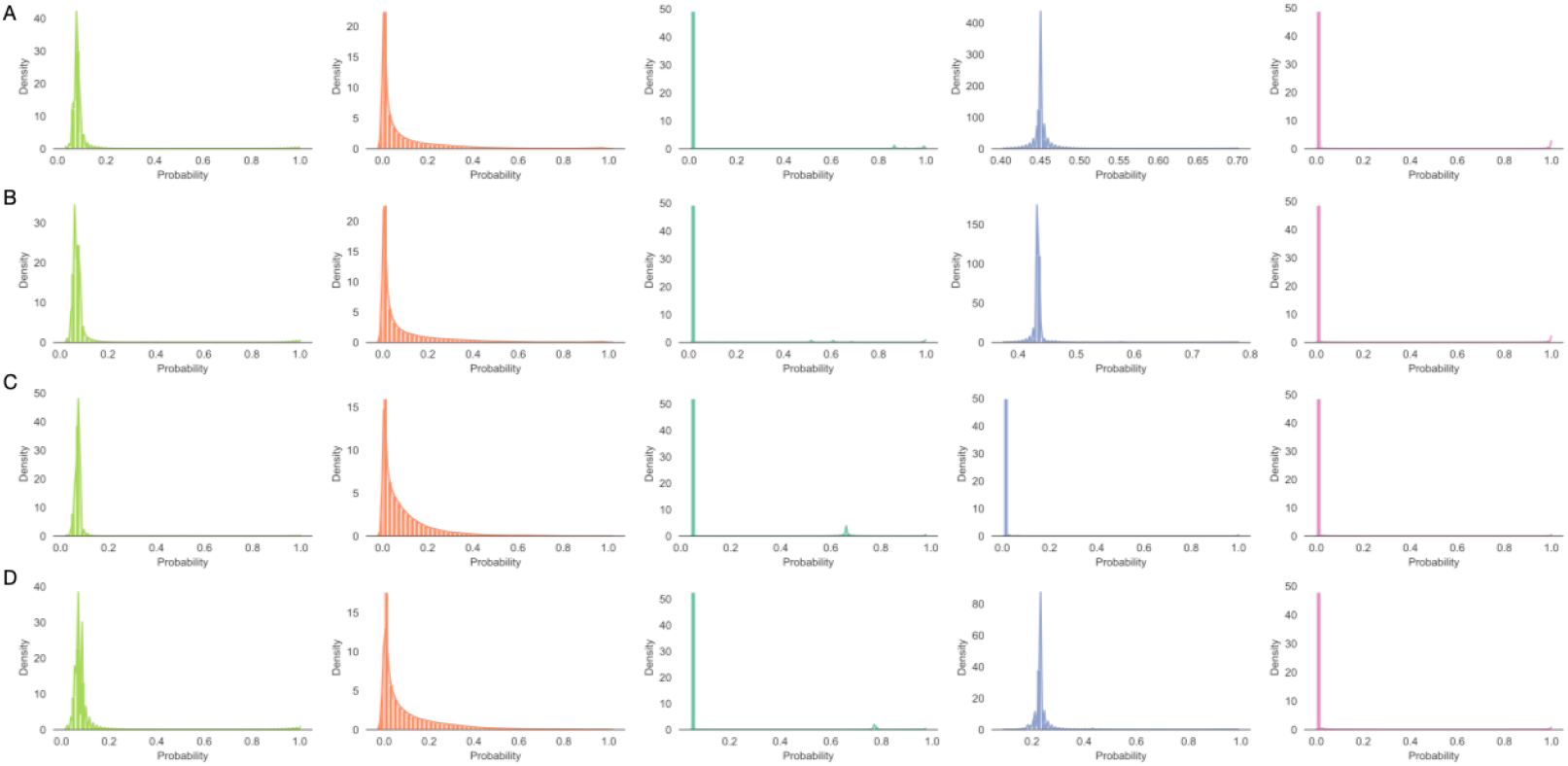
Model probability distributions assigned by machine learning algorithms, related to Figure 1. A. All Stroke, B. Ischemic Stroke, C. Sub-arachnoid Hemorrhage, D. Intracerebral Hemorrhage. First panel: Logistic Regression model with Elastic Net penalty probabiltiy distribution, Second panel: Random Forest probability distribution, Third panel: Logistic Regression model with L1 penalty probability distribution, Fourth panel: Adaboost probability distribution, Fifth panel: Gradient boosting probability distribution

**Figure A.2:**
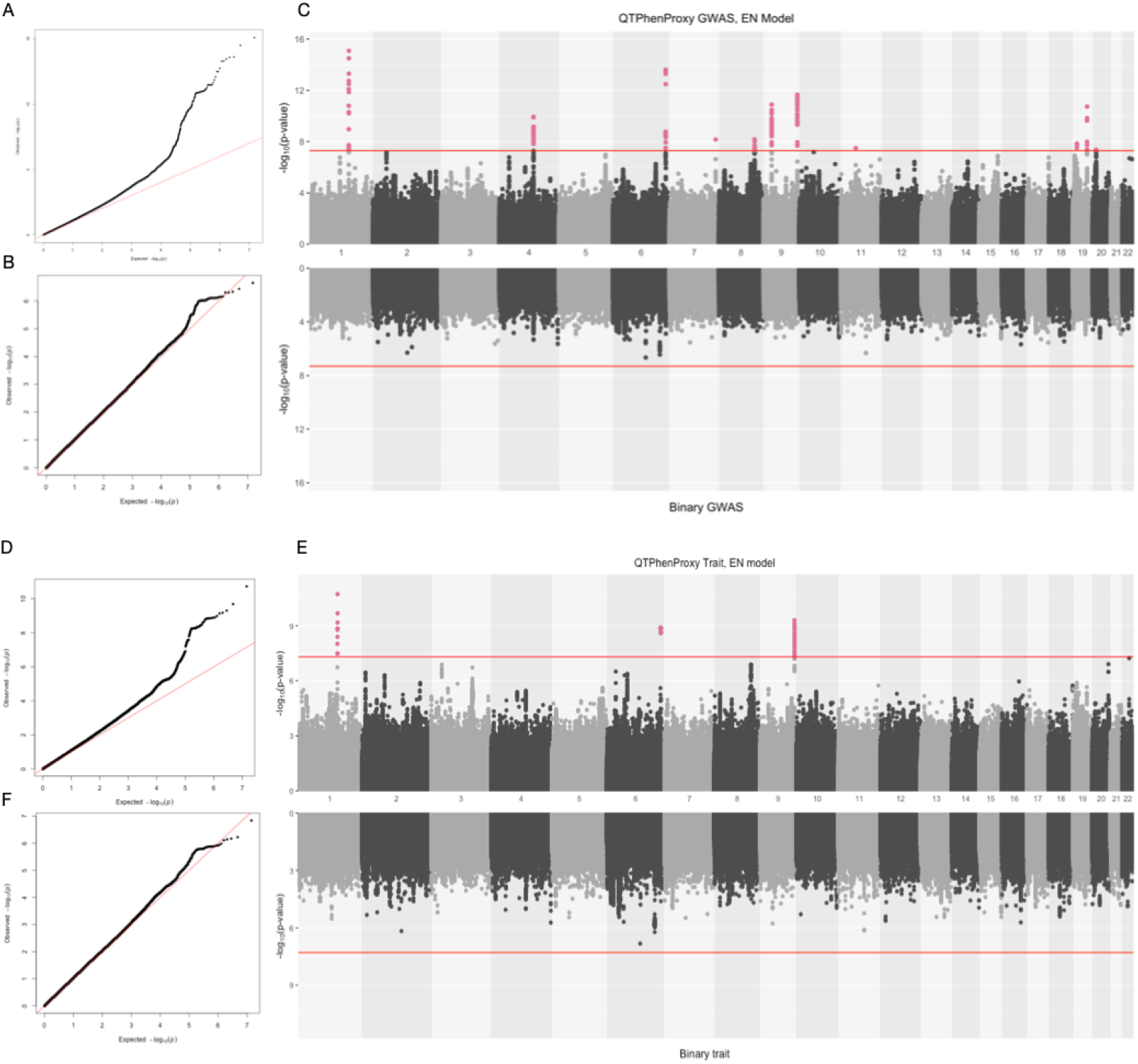
QQ Plots and Hudson plot of QTPhenProxy genome-wide association analysis, EN Model with Binary trait genome-wide association analysis for Ischemic Stroke, (Top) using QC1 quality control and (Bottom) using QC2 quality control, related to Figure 2. A. and D. Q-Q Plot for QTPhenProxy, EN Model GWAS. B.and E. Q-Q Plot for Binary trait GWAS. C. and F. Top manhattan plot is for QTPhenProxy, bottom plot for binary trait.Variants with p-value < 5e-08 are highlighted in pink, and the dashed lines are at the same value.

**Figure A.3:**
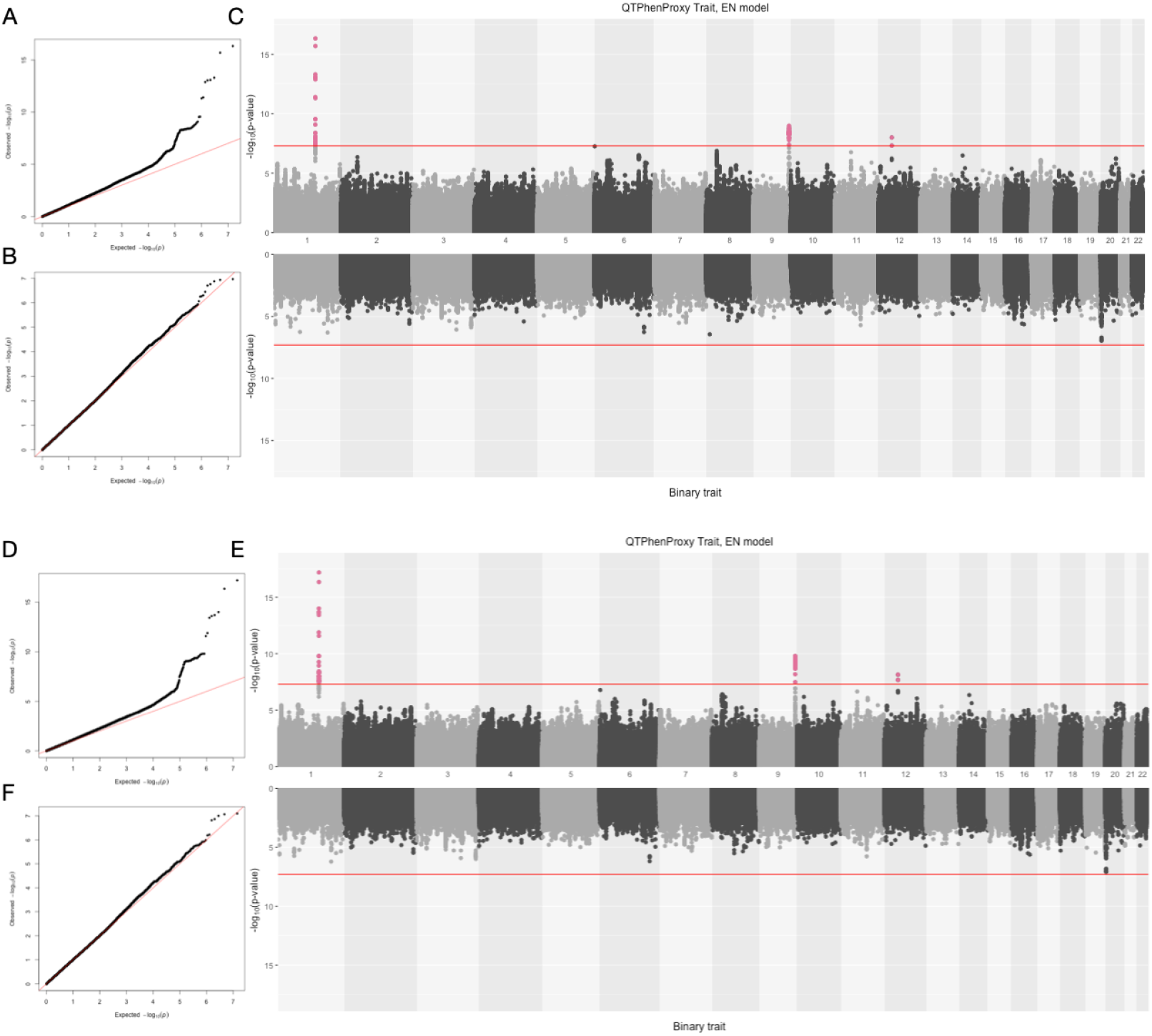
QQ Plots and Hudson plot of QTPhenProxy genome-wide association analysis, EN Model with Binary trait genome-wide association analysis for Subarachnoid Hemorrhage, (Top) using QC1 quality control and (Bottom) using QC2 quality control, related to Figure 2. A. and D. Q-Q Plot for QTPhenProxy, EN Model GWAS. B.and E. Q-Q Plot for Binary trait GWAS. C. and F. Top manhattan plot is for QTPhenProxy, bottom plot for binary trait.Variants with p-value < 5e-08 are highlighted in pink, and the dashed lines are at the same value.

**Figure A.4:**
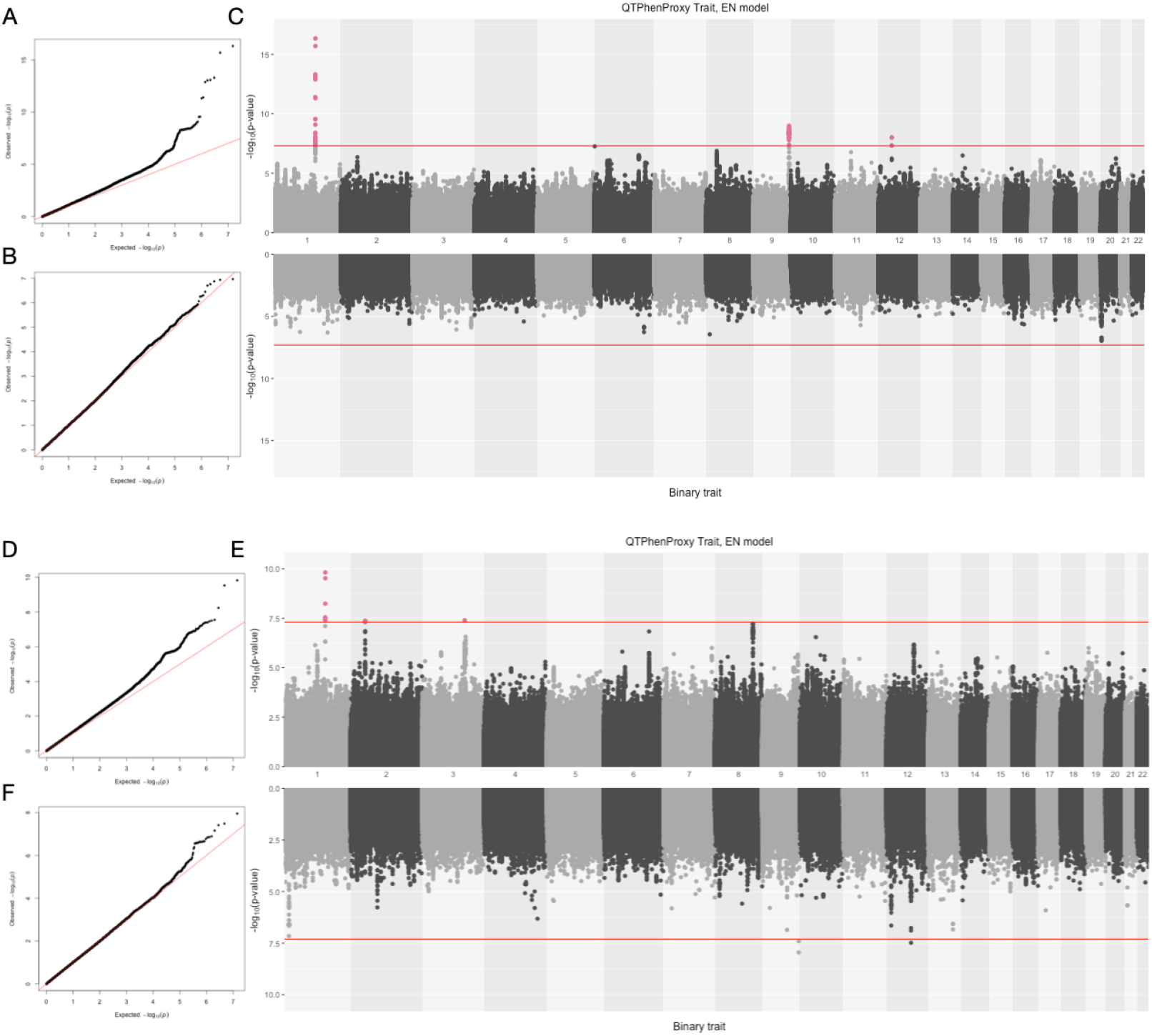
QQ Plots and Hudson plot of QTPhenProxy genome-wide association analysis, EN Model with Binary trait genome-wide association analysis for Intracerebral Hemorrhage, (Top) using QC1 quality control and (Bottom) using QC2 quality control, related to Figure 2. A. and D. Q-Q Plot for QTPhenProxy, EN Model GWAS. B.and E. Q-Q Plot for Binary trait GWAS. C. and F. Top manhattan plot is for QTPhenProxy, bottom plot for binary trait.Variants with p-value < 5e-08 are highlighted in pink, and the dashed lines are at the same value.

**Figure A.5:**
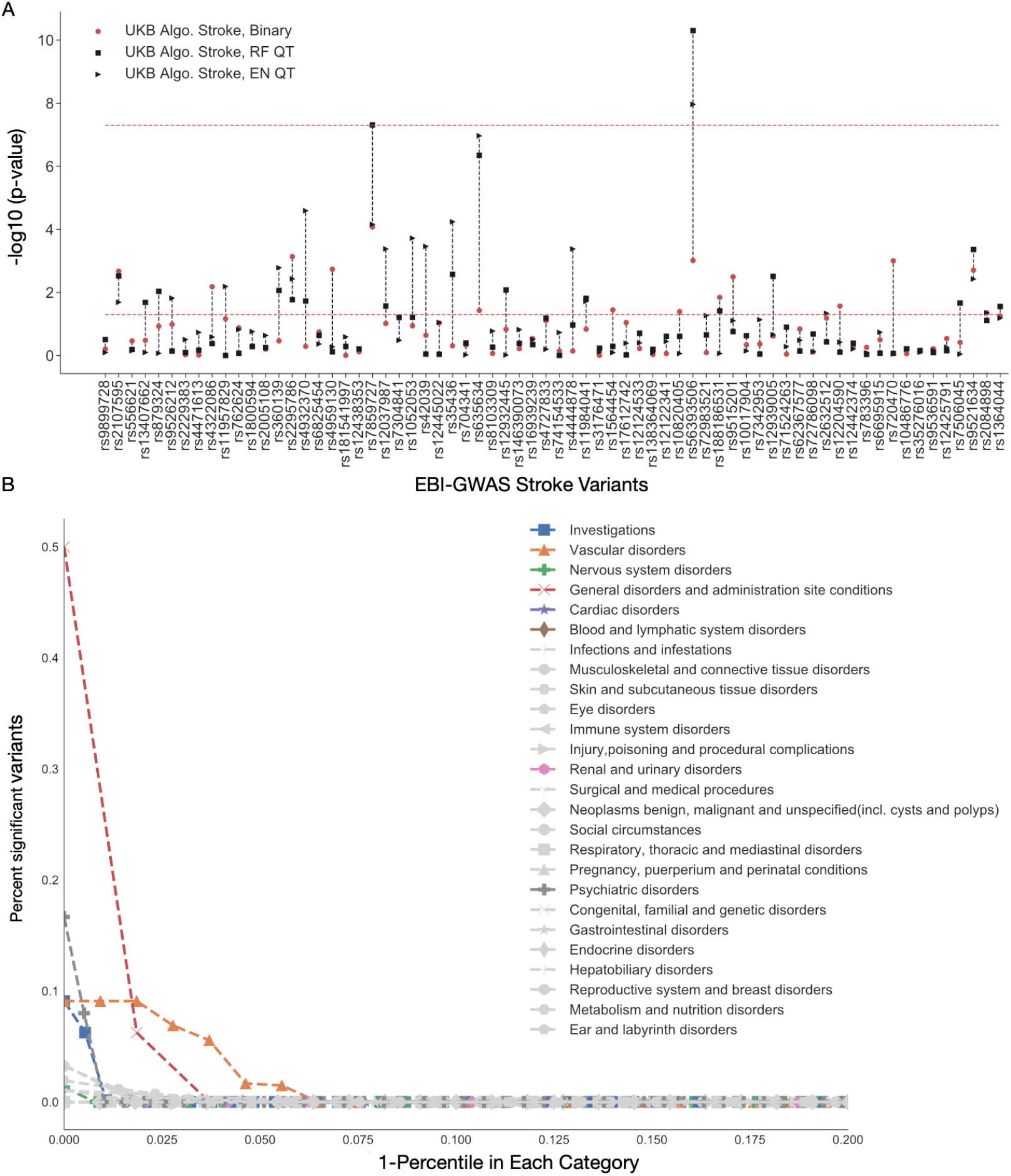
QTPhenProxy recovers known stroke variants and variants specific to related organ systems, related to Figure 3. Results from QTPhenProxy GWAS with QC2 quality control. (A) Horizontal axis shows Stroke variants catalogued in EBI-GWAS that have shown genomewide significance in previous studies. Black markers represent p-values of variants recovered by QTPhenProxy models (square=RF,Triangle=EN). (B) X-axis is the top percentile of marker sets in each category, Y-axis is the proportion of variants in marker sets that overlap with QTPhenProxy genome-wide significant variants. Each shape corresponds to a disease category. In color are top disease categories: Square=Investigations, Triangle=Vascular disorders, Cross=Nervous system disorders, X=General disorders and administration site conditions, Star=cardiac disorders, and Diamond=Blood and lymphatic system disorders.

**Figure A.6:**
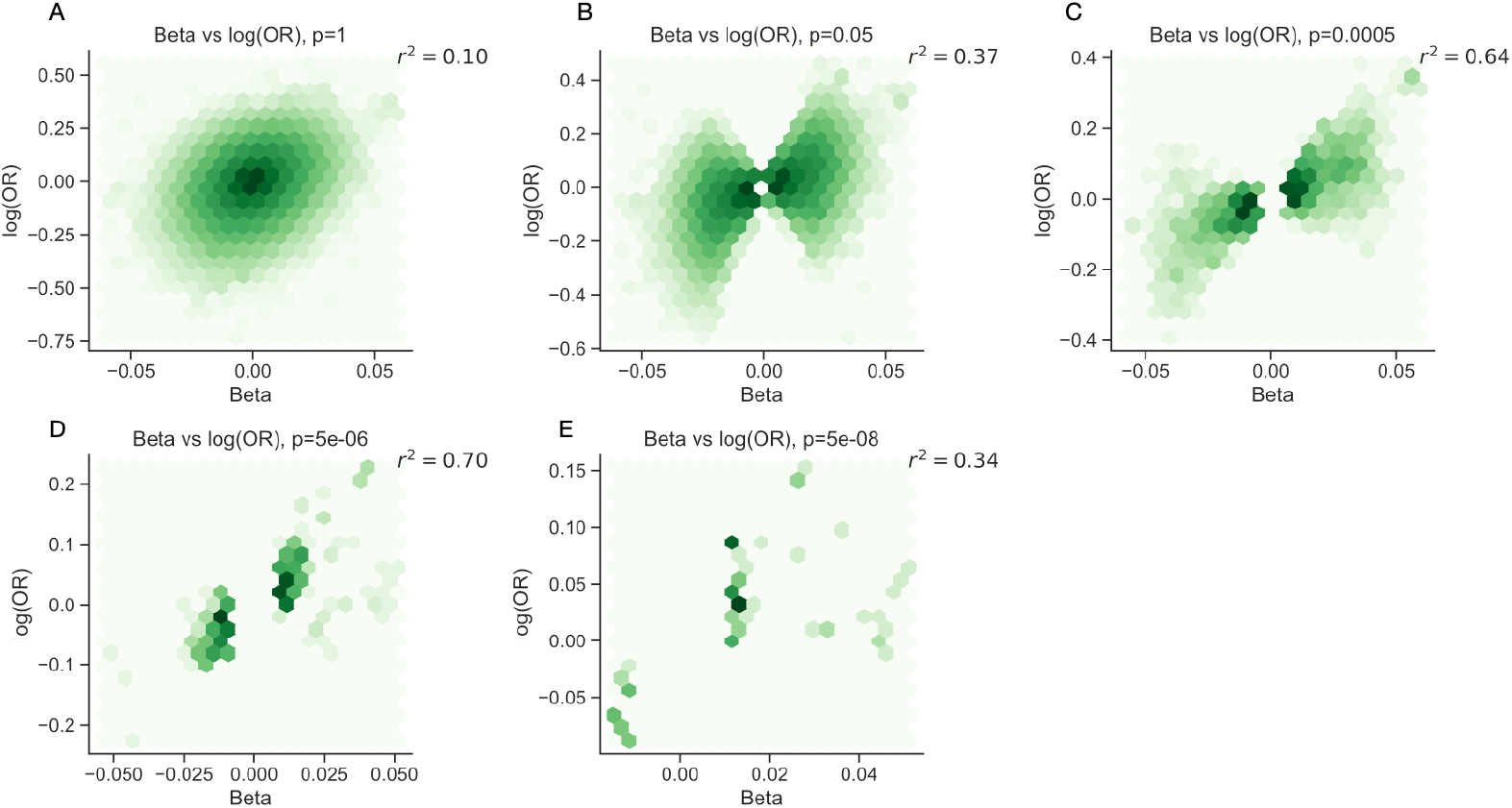
Correlation between beta and odds ratio effect sizes for variants at varying degrees of significance, related to Figure 4. (A-E) Correlation between QTPhenProxy, EN Model for Stroke GWAS and Binary trait GWAS effect size, QC1 quality control. X-axis is QTPhenProxy GWAS beta-coefficients, Y-axis is log base 10 of the Binary GWAS odds ratios. Pearson correlation is recorded in top right corner. From left to right, top to bottom, variants included decreases by restricting p-value.

**Figure A.7:**
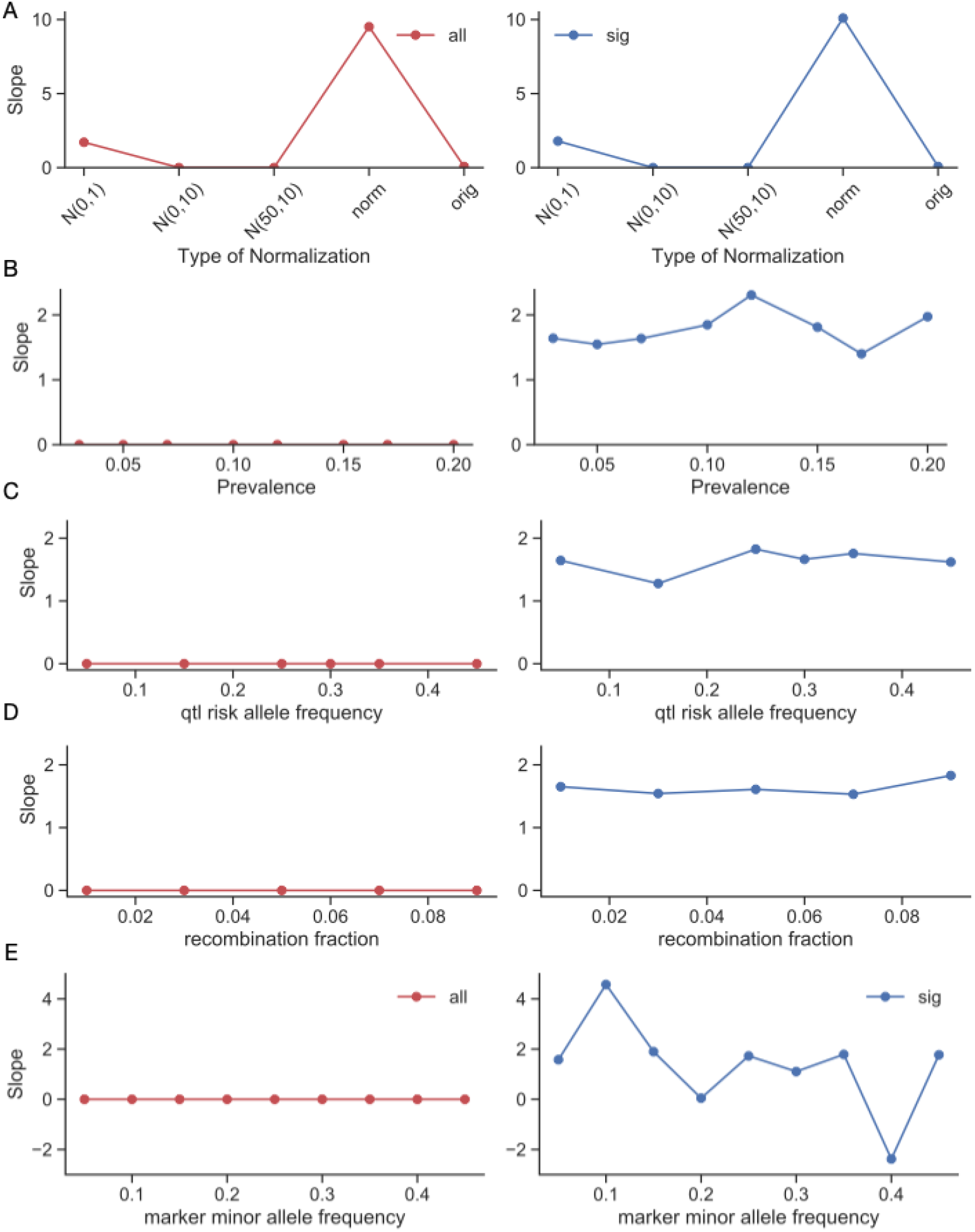
Slope of correlation between beta from simulated quantitative trait and log(odds ratio) of simulated binary trait with varying simulation parameters. Left panels calculate slope with all variants, right panels calculate slope with variants with p-value <0.005. (A) varies the transformation of the probability distribution, where *N(0,1)* is a normal distribution with mean 0, variance 1, *N(0,10)* is a normal distribution with mean 0, variance 10,*N(10,50)* is a normal distribution with mean 10, variance 50, *norm* is normalized by the maximum value, and *orig* is the original distribution. (B) varies the prevalence of the trait, (C) varies the causal allele frequency, (D) varies the recombination fraction between the causal allele and marker allele, and (E) varies the marker minor allele frequency.

## Appendix B. Supplementary Tables

**Table B.1:** Phenotyping Models Performance. *rf*: Random Forest, *ab*= Adaboost, *lr*=Logistic regression with L1 penalty, gb=Gradient Boosting, en=Logistic Regression with elastic net penalty models. *AUROC*: Area under the Receiver Operating Curve, *AUPRC*: Area under Precision-Recall curve. 95% confidence intervals shown in parentheses

| Disease | Model | AUROC | AUPRC | Maximum F1 |
| --- | --- | --- | --- | --- |
| Ischemic Stroke | rf | 0.953(0.000279) | 0.248(0.00201) | 0.326(0.00229) |
|  | ab | 0.950(0.000504) | 0.246(0.00465) | 0.316(0.00335) |
|  | lr | 0.946(0.000409) | 0.239(0.0021) | 0.322(0.00196) |
|  | gb | 0.952(0.000686) | 0.210(0.00479) | 0.294(0.00319) |
|  | en | 0.943(0.000362) | 0.209(0.00193) | 0.295(0.00199) |
| SAH Stroke | rf | 0.875(0.000746) | 0.144(0.00299) | 0.242(0.00423) |
|  | ab | 0.859(0.00186) | 0.171(0.0062) | 0.271(0.00807) |
|  | lr | 0.850(0.00115) | 0.151(0.00419) | 0.291(0.00509) |
|  | gb | 0.863(0.00149) | 0.113(0.00379) | 0.203(0.00513) |
|  | en | 0.863(0.000958) | 0.109(0.00377) | 0.198(0.00497) |
| ICH Stroke | rf | 0.903(0.000924) | 0.0483(0.00143) | 0.116(0.00253) |
|  | ab | 0.900(0.00171) | 0.0480(0.00196) | 0.114(0.00305) |
|  | lr | 0.879(0.00145) | 0.0270(0.000709) | 0.0762(0.00137) |
|  | gb | 0.900(0.00123) | 0.0365(0.00125) | 0.0935(0.00227) |
|  | en | 0.903(0.00118) | 0.0329(0.000755) | 0.0861(0.00146) |
| Stroke | rf | 0.899(0.000404) | 0.326(0.0016) | 0.375(0.00147) |
|  | ab | 0.906(0.000711) | 0.335(0.00423) | 0.380(0.00262) |
|  | lr | 0.905(0.000528) | 0.314(0.00143) | 0.368(0.00154) |
|  | gb | 0.903(0.00047) | 0.292(0.00245) | 0.355(0.00151) |
|  | en | 0.896(0.000448) | 0.282(0.00113) | 0.350(0.000962) |

**Table B.2:**
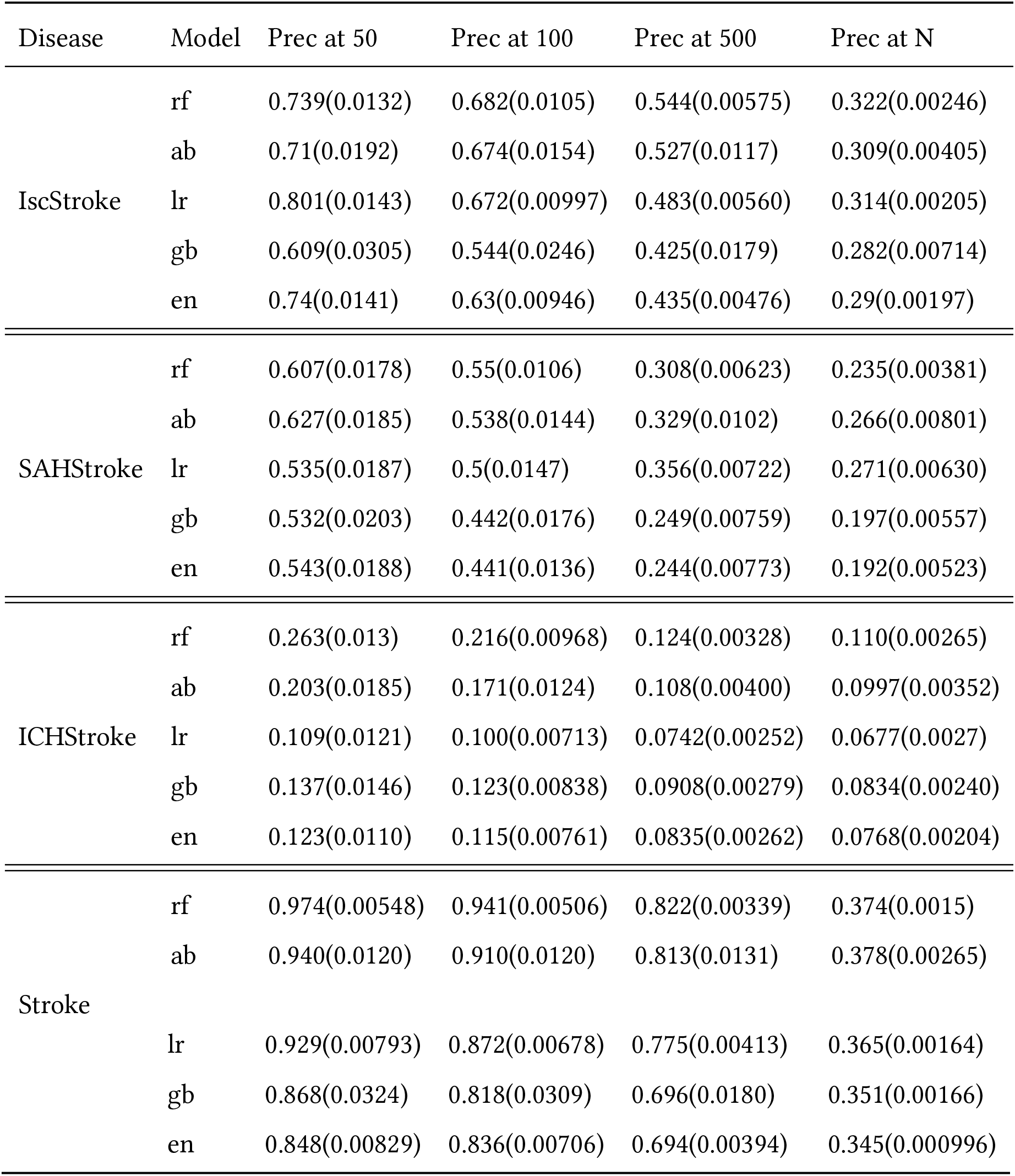
Precision at top 50, 100, 500, and N cases probabilities of phenotyping models. *rf*: Random Forest, *ab*= Adaboost, *lr*=Logistic regression with L1 penalty, gb=Gradient Boosting, en=Logistic Regression with elastic net penalty models. *Prec*: Precision. 95% confidence intervals

**Table B.3:** Sensitivity, Specificity, Positive Predictive Value, and Negative Predictive Value of phenotyping models. *rf*: Random Forest, *ab*= Adaboost, *lr*=Logistic regression with L1 penalty, gb=Gradient Boosting, en=Logistic Regression with elastic net penalty models. *PPV*: Positive Predictive Value, *NPV*: Negative Predictive Value. 95% confidence intervals shown in parentheses

| Disease | Model | Sensitivity | Specificity | PPV | NPV |
| --- | --- | --- | --- | --- | --- |
| Ischemic Stroke | rf | 0.347(0.00483) | 0.996(0.000133) | 0.00185(2.58e-05) | 0.997(2.52e-05) |
|  | ab | 0.369(0.00744) | 0.995(0.000256) | 0.00196(3.99e-05) | 0.997(3.86e-05) |
|  | lr | 0.362(0.00528) | 0.995(0.000127) | 0.00192(2.82e-05) | 0.997(2.76e-05) |
|  | gb | 0.368(0.00868) | 0.994(0.000354) | 0.00196(4.67e-05) | 0.997(4.50e-05) |
|  | en | 0.332(0.00624) | 0.995(0.000195) | 0.00177(3.35e-05) | 0.996(3.24e-05) |
| SAH Stroke | rf | 0.215(0.00503) | 0.999(5.47e-05) | 0.000354(8.28e-06) | 0.999(8.21e-06) |
|  | ab | 0.272(0.00924) | 0.999(7.14e-05) | 0.000447(1.52e-05) | 0.999(1.51e-05) |
|  | lr | 0.266(0.00764) | 0.999(6.34e-05) | 0.000437(1.26e-05) | 0.999(1.25e-05) |
|  | gb | 0.204(0.00598) | 0.999(9.43e-05) | 0.000336(9.86e-06) | 0.999(9.75e-06) |
|  | en | 0.201(0.00597) | 0.999(0.000126) | 0.000330(9.85e-06) | 0.999(9.68e-06) |
| ICH Stroke | rf | 0.133(0.00715) | 0.998(0.000140) | 0.000174(9.40e-06) | 0.999(9.22e-06) |
|  | ab | 0.167(0.0113) | 0.998(0.000219) | 0.000220(1.49e-05) | 0.999(1.47e-05) |
|  | lr | 0.177(0.0211) | 0.995(0.000742) | 0.000233(2.79e-05) | 0.999(2.69e-05) |
|  | gb | 0.141(0.00950) | 0.998(0.000268) | 0.000185(1.25e-05) | 0.999(1.22e-05) |
|  | en | 0.146(0.0123) | 0.997(0.000398) | 0.000192(1.62e-05) | 0.999(1.57e-05) |
| Stroke | rf | 0.377(0.00414) | 0.989(0.000286) | 0.00648(7.24e-05) | 0.989(6.74e-05) |
|  | ab | 0.390(0.00559) | 0.989(0.000379) | 0.00671(9.76e-05) | 0.990(9.17e-05) |
|  | lr | 0.399(0.00499) | 0.987(0.000408) | 0.00688(8.81e-05) | 0.990(8.1e-05) |
|  | gb | 0.388(0.00462) | 0.986(0.000373) | 0.00670(8.15e-05) | 0.989(7.51e-05) |
|  | en | 0.393(0.00425) | 0.985(0.000336) | 0.00678(7.51e-05) | 0.990(6.90e-05) |

**Table B.4:**
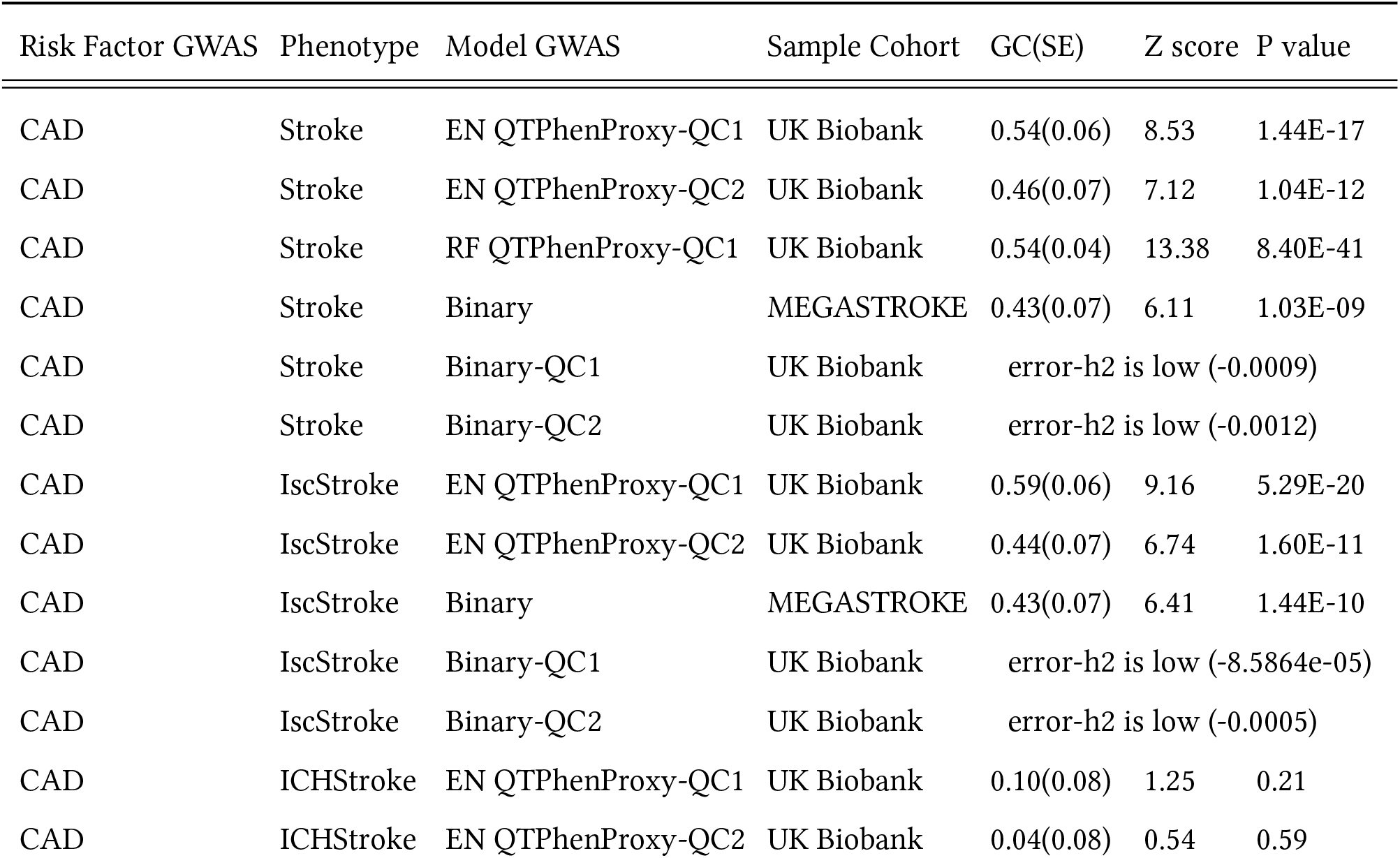
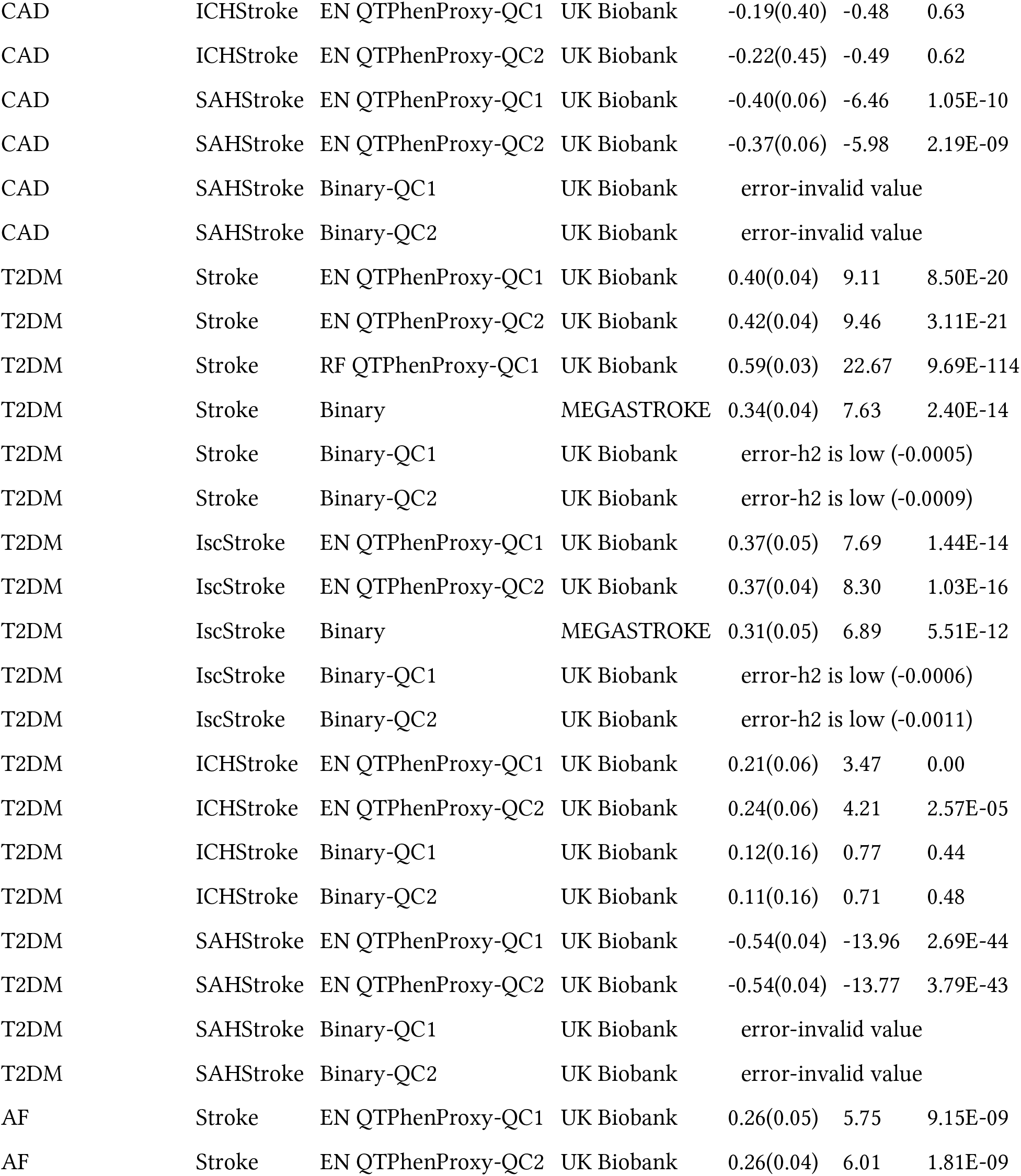
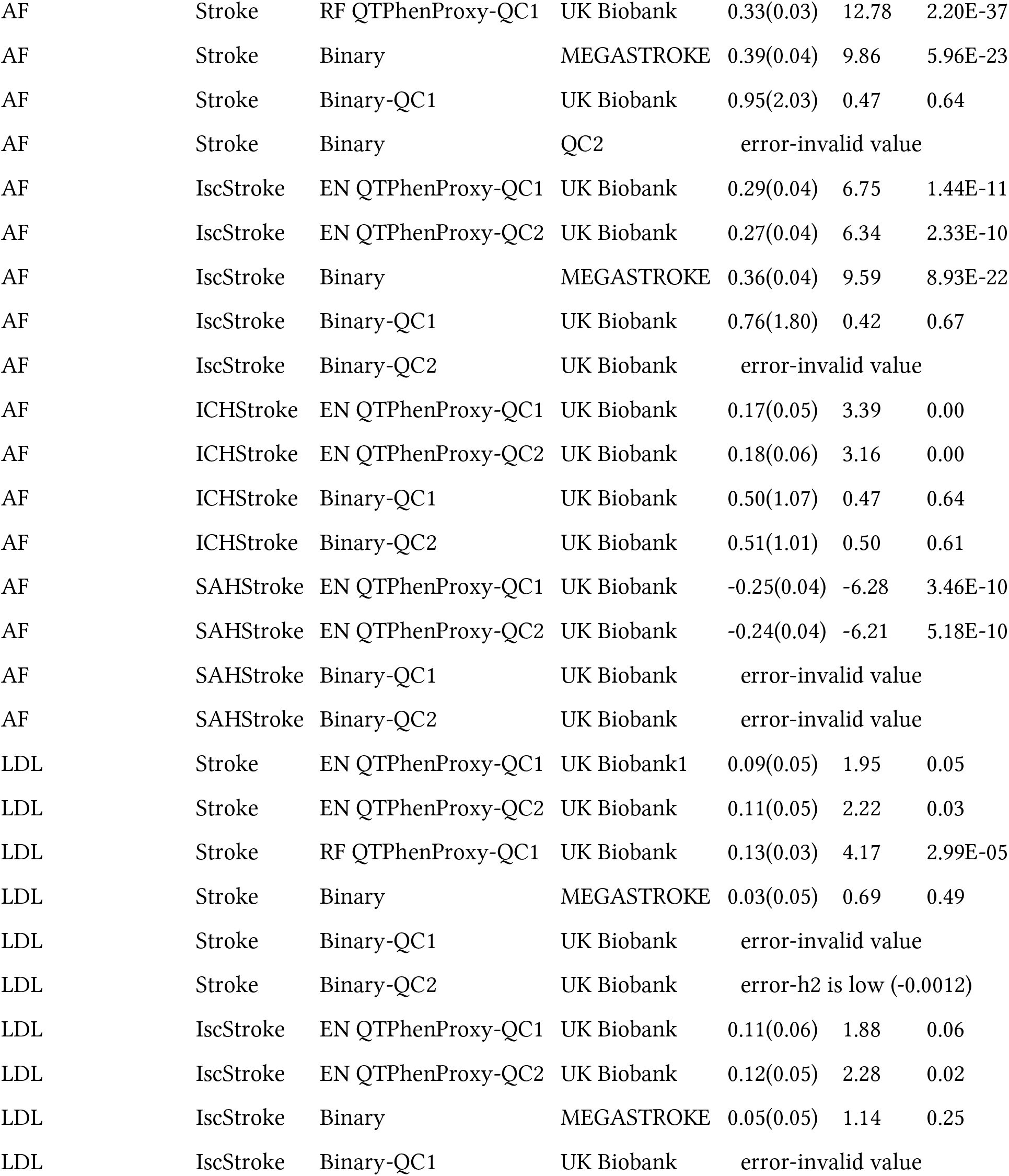
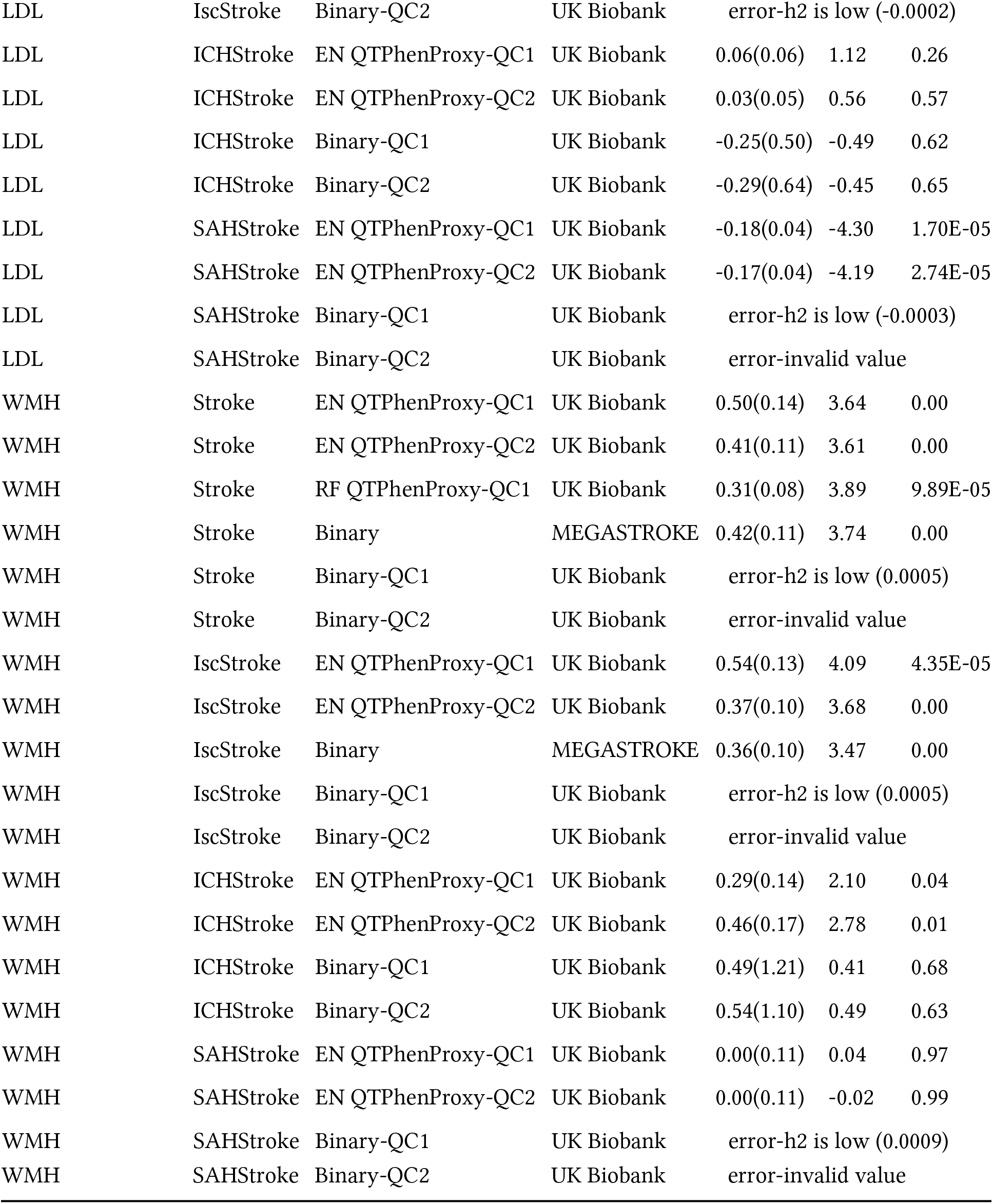
Genetic Correlation of Stroke Risk Factor GWAS with QTPhenProxy Models and MEGASTROKE. *GC*: Genetic Correlation, *SE*: Standard Error, *CAD*: Coronary Artery Disease, *T2DM*: Type 2 Diabetes Mellitus, *AF*: Atrial Fibrillation, *LDL*: Low-Density Lipoproteins *WMH*: White Matter Hyperintensities

| Risk Factor GWAS | Phenotype | Model GWAS | Sample Cohort | GC(SE) | Z score | P value |
| --- | --- | --- | --- | --- | --- | --- |
| CAD | Stroke | EN QTPhenProxy-QC1 | UK Biobank | 0.54(0.06) | 8.53 | 1.44E-17 |
| CAD | Stroke | EN QTPhenProxy-QC2 | UK Biobank | 0.46(0.07) | 7.12 | 1.04E-12 |
| CAD | Stroke | RF QTPhenProxy-QC1 | UK Biobank | 0.54(0.04) | 13.38 | 8.40E-41 |
| CAD | Stroke | Binary | MEGASTROKE | 0.43(0.07) | 6.11 | 1.03E-09 |
| CAD | Stroke | Binary-QC1 | UK Biobank | error-h2 is low (-0.0009) |  |  |
| CAD | Stroke | Binary-QC2 | UK Biobank | error-h2 is low (-0.0012) |  |  |
| CAD | IscStroke | EN QTPhenProxy-QC1 | UK Biobank | 0.59(0.06) | 9.16 | 5.29E-20 |
| CAD | IscStroke | EN QTPhenProxy-QC2 | UK Biobank | 0.44(0.07) | 6.74 | 1.60E-11 |
| CAD | IscStroke | Binary | MEGASTROKE | 0.43(0.07) | 6.41 | 1.44E-10 |
| CAD | IscStroke | Binary-QC1 | UK Biobank | error-h2 is low (-8.5864e-05) |  |  |
| CAD | IscStroke | Binary-QC2 | UK Biobank | error-h2 is low (-0.0005) |  |  |
| CAD | ICHStroke | EN QTPhenProxy-QC1 | UK Biobank | 0.10(0.08) | 1.25 | 0.21 |
| CAD | ICHStroke | EN QTPhenProxy-QC2 | UK Biobank | 0.04(0.08) | 0.54 | 0.59 |
| CAD | ICHStroke | EN QTPhenProxy-QC1 | UK Biobank | -0.19(0.40) | -0.48 | 0.63 |
| CAD | ICHStroke | EN QTPhenProxy-QC2 | UK Biobank | -0.22(0.45) | -0.49 | 0.62 |
| CAD | SAHStroke | EN QTPhenProxy-QC1 | UK Biobank | -0.40(0.06) | -6.46 | 1.05E-10 |
| CAD | SAHStroke | EN QTPhenProxy-QC2 | UK Biobank | -0.37(0.06) | -5.98 | 2.19E-09 |
| CAD | SAHStroke | Binary-QC1 | UK Biobank | error-invalid value |  |  |
| CAD | SAHStroke | Binary-QC2 | UK Biobank | error-invalid value |  |  |
| T2DM | Stroke | EN QTPhenProxy-QC1 | UK Biobank | 0.40(0.04) | 9.11 | 8.50E-20 |
| T2DM | Stroke | EN QTPhenProxy-QC2 | UK Biobank | 0.42(0.04) | 9.46 | 3.11E-21 |
| T2DM | Stroke | RF QTPhenProxy-QC1 | UK Biobank | 0.59(0.03) | 22.67 | 9.69E-114 |
| T2DM | Stroke | Binary | MEGASTROKE | 0.34(0.04) | 7.63 | 2.40E-14 |
| T2DM | Stroke | Binary-QC1 | UK Biobank | error-h2 is low (-0.0005) |  |  |
| T2DM | Stroke | Binary-QC2 | UK Biobank | error-h2 is low (-0.0009) |  |  |
| T2DM | IscStroke | EN QTPhenProxy-QC1 | UK Biobank | 0.37(0.05) | 7.69 | 1.44E-14 |
| T2DM | IscStroke | EN QTPhenProxy-QC2 | UK Biobank | 0.37(0.04) | 8.30 | 1.03E-16 |
| T2DM | IscStroke | Binary | MEGASTROKE | 0.31(0.05) | 6.89 | 5.51E-12 |
| T2DM | IscStroke | Binary-QC1 | UK Biobank | error-h2 is low (-0.0006) |  |  |
| T2DM | IscStroke | Binary-QC2 | UK Biobank | error-h2 is low (-0.0011) |  |  |
| T2DM | ICHStroke | EN QTPhenProxy-QC1 | UK Biobank | 0.21(0.06) | 3.47 | 0.00 |
| T2DM | ICHStroke | EN QTPhenProxy-QC2 | UK Biobank | 0.24(0.06) | 4.21 | 2.57E-05 |
| T2DM | ICHStroke | Binary-QC1 | UK Biobank | 0.12(0.16) | 0.77 | 0.44 |
| T2DM | ICHStroke | Binary-QC2 | UK Biobank | 0.11(0.16) | 0.71 | 0.48 |
| T2DM | SAHStroke | EN QTPhenProxy-QC1 | UK Biobank | -0.54(0.04) | -13.96 | 2.69E-44 |
| T2DM | SAHStroke | EN QTPhenProxy-QC2 | UK Biobank | -0.54(0.04) | -13.77 | 3.79E-43 |
| T2DM | SAHStroke | Binary-QC1 | UK Biobank | error-invalid value |  |  |
| T2DM | SAHStroke | Binary-QC2 | UK Biobank | error-invalid value |  |  |
| AF | Stroke | EN QTPhenProxy-QC1 | UK Biobank | 0.26(0.05) | 5.75 | 9.15E-09 |
| AF | Stroke | EN QTPhenProxy-QC2 | UK Biobank | 0.26(0.04) | 6.01 | 1.81E-09 |
| AF | Stroke | RF QTPhenProxy-QC1 | UK Biobank | 0.33(0.03) | 12.78 | 2.20E-37 |
| AF | Stroke | Binary | MEGASTROKE | 0.39(0.04) | 9.86 | 5.96E-23 |
| AF | Stroke | Binary-QC1 | UK Biobank | 0.95(2.03) | 0.47 | 0.64 |
| AF | Stroke | Binary | QC2 | error-invalid value |  |  |
| AF | IscStroke | EN QTPhenProxy-QC1 | UK Biobank | 0.29(0.04) | 6.75 | 1.44E-11 |
| AF | IscStroke | EN QTPhenProxy-QC2 | UK Biobank | 0.27(0.04) | 6.34 | 2.33E-10 |
| AF | IscStroke | Binary | MEGASTROKE | 0.36(0.04) | 9.59 | 8.93E-22 |
| AF | IscStroke | Binary-QC1 | UK Biobank | 0.76(1.80) | 0.42 | 0.67 |
| AF | IscStroke | Binary-QC2 | UK Biobank | error-invalid value |  |  |
| AF | ICHStroke | EN QTPhenProxy-QC1 | UK Biobank | 0.17(0.05) | 3.39 | 0.00 |
| AF | ICHStroke | EN QTPhenProxy-QC2 | UK Biobank | 0.18(0.06) | 3.16 | 0.00 |
| AF | ICHStroke | Binary-QC1 | UK Biobank | 0.50(1.07) | 0.47 | 0.64 |
| AF | ICHStroke | Binary-QC2 | UK Biobank | 0.51(1.01) | 0.50 | 0.61 |
| AF | SAHStroke | EN QTPhenProxy-QC1 | UK Biobank | -0.25(0.04) | -6.28 | 3.46E-10 |
| AF | SAHStroke | EN QTPhenProxy-QC2 | UK Biobank | -0.24(0.04) | -6.21 | 5.18E-10 |
| AF | SAHStroke | Binary-QC1 | UK Biobank | error-invalid value |  |  |
| AF | SAHStroke | Binary-QC2 | UK Biobank | error-invalid value |  |  |
| LDL | Stroke | EN QTPhenProxy-QC1 | UK Biobank1 | 0.09(0.05) | 1.95 | 0.05 |
| LDL | Stroke | EN QTPhenProxy-QC2 | UK Biobank | 0.11(0.05) | 2.22 | 0.03 |
| LDL | Stroke | RF QTPhenProxy-QC1 | UK Biobank | 0.13(0.03) | 4.17 | 2.99E-05 |
| LDL | Stroke | Binary | MEGASTROKE | 0.03(0.05) | 0.69 | 0.49 |
| LDL | Stroke | Binary-QC1 | UK Biobank | error-invalid value |  |  |
| LDL | Stroke | Binary-QC2 | UK Biobank | error-h2 is low (-0.0012) |  |  |
| LDL | IscStroke | EN QTPhenProxy-QC1 | UK Biobank | 0.11(0.06) | 1.88 | 0.06 |
| LDL | IscStroke | EN QTPhenProxy-QC2 | UK Biobank | 0.12(0.05) | 2.28 | 0.02 |
| LDL | IscStroke | Binary | MEGASTROKE | 0.05(0.05) | 1.14 | 0.25 |
| LDL | IscStroke | Binary-QC1 | UK Biobank | error-invalid value |  |  |
| LDL | IscStroke | Binary-QC2 | UK Biobank | error-h2 is low (-0.0002) |  |  |
| LDL | ICHStroke | EN QTPhenProxy-QC1 | UK Biobank | 0.06(0.06) | 1.12 | 0.26 |
| LDL | ICHStroke | EN QTPhenProxy-QC2 | UK Biobank | 0.03(0.05) | 0.56 | 0.57 |
| LDL | ICHStroke | Binary-QC1 | UK Biobank | -0.25(0.50) | -0.49 | 0.62 |
| LDL | ICHStroke | Binary-QC2 | UK Biobank | -0.29(0.64) | -0.45 | 0.65 |
| LDL | SAHStroke | EN QTPhenProxy-QC1 | UK Biobank | -0.18(0.04) | -4.30 | 1.70E-05 |
| LDL | SAHStroke | EN QTPhenProxy-QC2 | UK Biobank | -0.17(0.04) | -4.19 | 2.74E-05 |
| LDL | SAHStroke | Binary-QC1 | UK Biobank | error-h2 is low (-0.0003) |  |  |
| LDL | SAHStroke | Binary-QC2 | UK Biobank | error-invalid value |  |  |
| WMH | Stroke | EN QTPhenProxy-QC1 | UK Biobank | 0.50(0.14) | 3.64 | 0.00 |
| WMH | Stroke | EN QTPhenProxy-QC2 | UK Biobank | 0.41(0.11) | 3.61 | 0.00 |
| WMH | Stroke | RF QTPhenProxy-QC1 | UK Biobank | 0.31(0.08) | 3.89 | 9.89E-05 |
| WMH | Stroke | Binary | MEGASTROKE | 0.42(0.11) | 3.74 | 0.00 |
| WMH | Stroke | Binary-QC1 | UK Biobank | error-h2 is low (0.0005) |  |  |
| WMH | Stroke | Binary-QC2 | UK Biobank | error-invalid value |  |  |
| WMH | IscStroke | EN QTPhenProxy-QC1 | UK Biobank | 0.54(0.13) | 4.09 | 4.35E-05 |
| WMH | IscStroke | EN QTPhenProxy-QC2 | UK Biobank | 0.37(0.10) | 3.68 | 0.00 |
| WMH | IscStroke | Binary | MEGASTROKE | 0.36(0.10) | 3.47 | 0.00 |
| WMH | IscStroke | Binary-QC1 | UK Biobank | error-h2 is low (0.0005) |  |  |
| WMH | IscStroke | Binary-QC2 | UK Biobank | error-invalid value |  |  |
| WMH | ICHStroke | EN QTPhenProxy-QC1 | UK Biobank | 0.29(0.14) | 2.10 | 0.04 |
| WMH | ICHStroke | EN QTPhenProxy-QC2 | UK Biobank | 0.46(0.17) | 2.78 | 0.01 |
| WMH | ICHStroke | Binary-QC1 | UK Biobank | 0.49(1.21) | 0.41 | 0.68 |
| WMH | ICHStroke | Binary-QC2 | UK Biobank | 0.54(1.10) | 0.49 | 0.63 |
| WMH | SAHStroke | EN QTPhenProxy-QC1 | UK Biobank | 0.00(0.11) | 0.04 | 0.97 |
| WMH | SAHStroke | EN QTPhenProxy-QC2 | UK Biobank | 0.00(0.11) | -0.02 | 0.99 |
| WMH | SAHStroke | Binary-QC1 | UK Biobank | error-h2 is low (0.0009) |  |  |

Table B.4: **Genetic Correlation of Stroke Risk Factor GWAS with QTPhenProxy Models and MEGASTROKE.**
|  |  |  |  |  |
| --- | --- | --- | --- | --- |
| WMH | SAHStroke | Binary-QC2 | UK Biobank | error-invalid value |

### Appendix B.1. Supplementary Information Table Titles

1. LD Hub GC correlation results of QTPhenProxy Stroke EN Model, QC1 quality control.
2. LD Hub GC correlation results of QTPhenProxy Stroke EN Model, QC2 quality control.
3. GWAS resources for Supplementary Table B.4.

### Appendix B.2. Data and Code Availability Statements

The EBI-GWAS marker set-MedDRA mappings and code generated in this study will be made publicly available online at https://github.com/pthangaraj/QTPhenProxy.

---

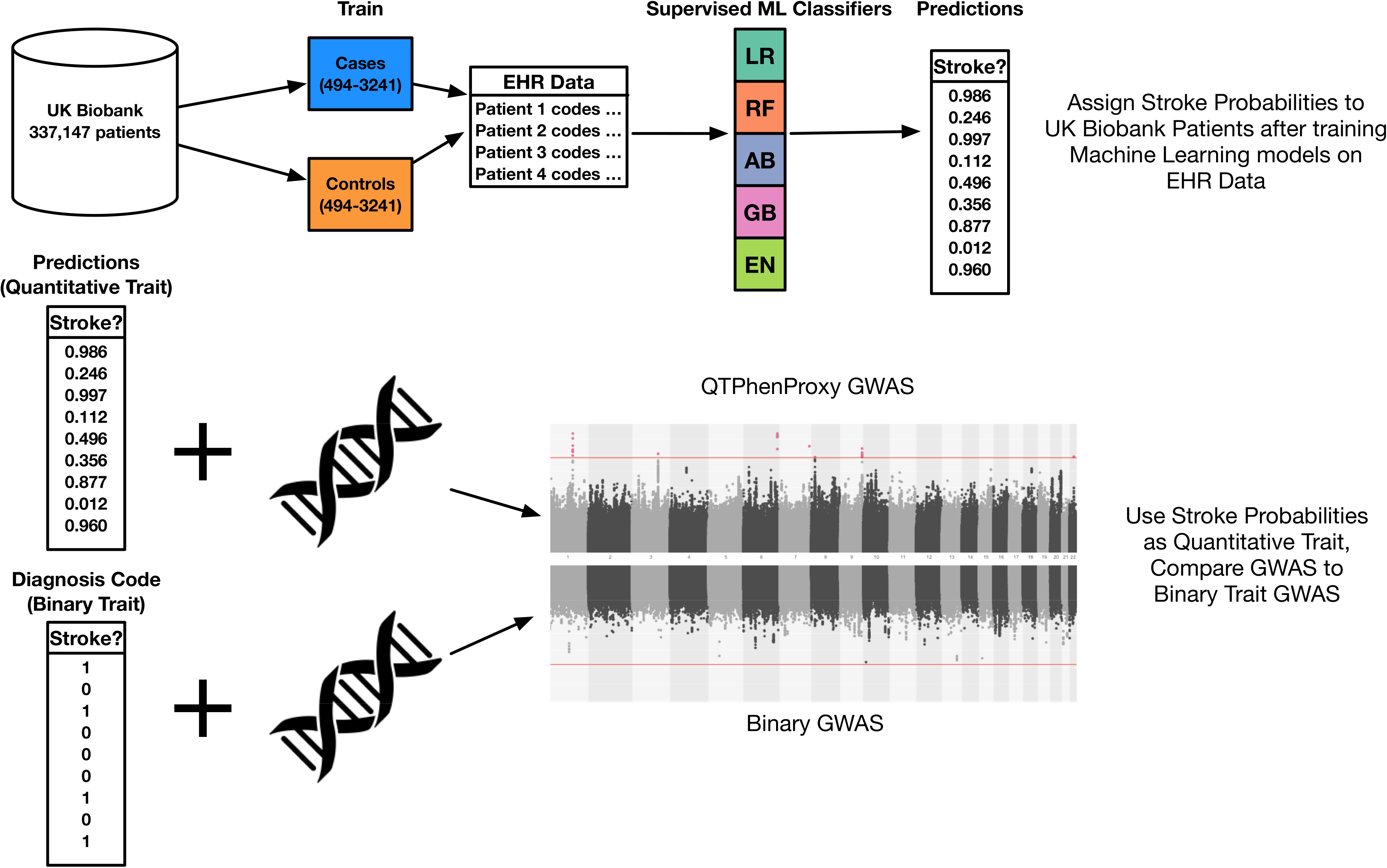

## Notes

### Competing Interest Statement

The authors have declared no competing interest.

## References

Abraham, G., Malik, R., Yonova-Doing, E., Salim, A., Wang, T., Danesh, J., Butterworth, A.S., Howson, J.M.M., Inouye, M., Dichgans, M., 2019. Genomic risk score offers predictive performance comparable to clinical risk factors for ischaemic stroke. Nature Communications 10, 5819. URL: http://www.nature.com/articles/s41467-019-13848-1, doi:10.1038/s41467-019-13848-1.

Almasy, L., Blangero, J., 1998. Multipoint Quantitative-Trait Linkage Analysis in General Pedigrees. The American Journal of Human Genetics 62, 1198–1211. URL: https://linkinghub.elsevier.com/retrieve/pii/S0002929707615420, doi:10.1086/301844.

Asselta, R., Peyvandi, F., 2009. Factor V deficiency. Seminars in Thrombosis and Hemostasis 35, 382–389. doi: 10.1055/s-0029-1225760.

Band, G., Marchini, J., 2018. BGEN: a binary file format for imputed genotype and haplotype data. bioRxiv URL: https://www.biorxiv.org/content/10.1101/308296v2, doi:10.1101/308296.

Brown, E., 1999. Coding of Data, MedDRA and other Medical Technologies, in: Clinical Data Management. Chichester, UK, p. 177. URL: 10.1002/0470846364.ch10.

Brown, M.S., Goldstein, J.L., 1979. Receptor-mediated endocytosis: insights from the lipoprotein receptor system. Proceedings of the National Academy of Sciences of the United States of America 76, 3330–3337.

Bulik-Sullivan, B.K., Loh, P.R., Finucane, H.K., Ripke, S., Yang, J., Patterson, N., Daly, M.J., Price, A.L., Neale, B.M., 2015. LD Score regression distinguishes confounding from polygenicity in genome-wide association studies. Nature Genetics 47, 291–295. URL: https://www.nature.com/articles/ng.3211, doi:10.1038/ng.3211.

Buniello, A., MacArthur, J.A.L., Cerezo, M., Harris, L.W., Hayhurst, J., Malangone, C., McMahon, A., Morales, J., Mountjoy, E., Sollis, E., Suveges, D., Vrousgou, O., Whetzel, P.L., Amode, R., Guillen, J.A., Riat, H.S., Trevanion, S.J., Hall, P., Junkins, H., Flicek, P., Burdett, T., Hindorff, L.A., Cunningham, F., Parkinson, H., 2019. The NHGRI-EBI GWAS Catalog of published genome-wide association studies, targeted arrays and summary statistics 2019. Nucleic Acids Research 47, D1005–D1012. URL: https://academic.oup.com/nar/article/47/D1/D1005/5184712, doi:10.1093/nar/gky1120.

Bycroft, C., Freeman, C., Petkova, D., Band, G., Elliott, L.T., Sharp, K., Motyer, A., Vukcevic, D., Delaneau, O., O’Connell, J., Cortes, A., Welsh, S., Young, A., Effingham, M., McVean, G., Leslie, S., Allen, N., Donnelly, P., Marchini, J., 2018. The UK Biobank resource with deep phenotyping and genomic data. Nature 562, 203–209. URL: http://www.nature.com/articles/s41586-018-0579-z, doi:10.1038/s41586-018-0579-z.

Chang, C.C., Chow, C.C., Tellier, L.C., Vattikuti, S., Purcell, S.M., Lee, J.J., 2015. Second-generation PLINK: rising to the challenge of larger and richer datasets. GigaScience 4, 7. URL: https://academic.oup.com/gigascience/article-lookup/doi/10.1186/s13742-015-0047-8, doi:10.1186/s13742-015-0047-8.

DeBoever, C., Tanigawa, Y., Aguirre, M., McInnes, G., Lavertu, A., Rivas, M.A., 2019. Assessing digital phenotyping to enhance genetic studies of human diseases. preprint. Genetics. URL: http://biorxiv.org/lookup/doi/10.1101/738856, doi:10.1101/738856.

Erdmann, J., Grosshennig, A., Braund, P.S., Konig, I.R., Hengstenberg, C., Hall, A.S., Linsel-Nitschke, P., Kathiresan, S., Wright, B., Tregouet, D.A., Cambien, F., Bruse, P., Aherrahrou, Z., Wagner, A.K., Stark, K., Schwartz, S.M., Salomaa, V., Elosua, R., Melander, O., Voight, B.F., O’Donnell, C.J., Peltonen, L., Siscovick, D.S., Altshuler, D., Merlini, P.A., Peyvandi, F., Bernardinelli, L., Ardissino, D., Schillert, A., Blankenberg, S., Zeller, T., Wild, P., Schwarz, D.F., Tiret, L., Perret, C., Schreiber, S., El Mokhtari, N.E., Schafer, A., Marz, W., Renner, W., Bugert, P., KlÃŒter, H., Schrezenmeir, J., Rubin, D., Ball, S.G., Balmforth, A.J., Wichmann, H.E., Meitinger, T., Fischer, M., Meisinger, C., Baumert, J., Peters, A., Ouwehand, W.H., Italian Atherosclerosis, Thrombosis, and Vascular Biology Working Group, Myocardial Infarction Genetics Consortium, Wellcome Trust Case Control Consortium, Cardiogenics Consortium, Deloukas, P., Thompson, J.R., Ziegler, A., Samani, N.J., Schunkert, H., 2009. New susceptibility locus for coronary artery disease on chromosome 3q22.3. Nature Genetics 41, 280–282.

Erqou, S., Thompson, A., Angelantonio, E.D., Saleheen, D., Kaptoge, S., Marcovina, S., Danesh, J., 2010. Apolipoprotein(a) Isoforms and the Risk of Vascular Disease: Systematic Review of 40 Studies Involving 58,000 Participants. Journal of the American College of Cardiology 55, 2160–2167. URL: http://www.onlinejacc.org/content/55/19/2160, doi:10.1016/j.jacc.2009.10.080.

Farah, C., Michel, L.Y.M., Balligand, J.L., 2018. Nitric oxide signalling in cardiovascular health and disease. Nature Reviews Cardiology 15, 292–316. URL: https://www.nature.com/articles/nrcardio.2017.224, doi:10.1038/nrcardio.2017.224.

Fessler, M.B., Arndt, P.G., Frasch, S.C., Lieber, J.G., Johnson, C.A., Murphy, R.C., Nick, J.A., Bratton, D.L., Malcolm, K.C., Worthen, G.S., 2004. Lipid Rafts Regulate Lipopolysaccharide-induced Activation of Cdc42 and Inflammatory Functions of the Human Neutrophil. Journal of Biological Chemistry 279, 39989–39998. URL: http://www.jbc.org/lookup/doi/10.1074/jbc.M401080200, doi:10.1074/jbc.M401080200.

Gao Yang, Stuart Deborah, Takahishi Takamune, Kohan Donald E., 2018. NephronâSpecific Disruption of Nitric Oxide Synthase 3 Causes Hypertension and Impaired Salt Excretion. Journal of the American Heart Association 7, e009236. URL: https://www.ahajournals.org/doi/full/10.1161/JAHA.118.009236, doi:10.1161/JAHA.118.009236.

Hashimoto, K., Ochi, H., Sunamura, S., Kosaka, N., Mabuchi, Y., Fukuda, T., Yao, K., Kanda, H., Ae, K., Okawa, A., Akazawa, C., Ochiya, T., Futakuchi, M., Takeda, S., Sato, S., 2018. Cancer-secreted hsa-miR-940 induces an osteoblastic phenotype in the bone metastatic microenvironment via targeting ARHGAP1 and FAM134a. Proceedings of the National Academy of Sciences 115, 2204–2209. URL: https://www.pnas.org/content/115/9/2204, doi:10.1073/pnas.1717363115.

Healthcare Cost and Utilization Project,. Multi-level procedures ccs categories. https://www.hcup-us.ahrq.gov/toolssoftware/ccs10/ccs10.jsp#download.

Hinds, D.A., Buil, A., Ziemek, D., Martinez-Perez, A., Malik, R., Folkersen, L., Germain, M., Malarstig, A., Brown, A., Soria, J.M., Dichgans, M., Bing, N., Franco-Cereceda, A., Souto, J.C., Dermitzakis, E.T., Hamsten, A., Worrall, B.B., Tung, J.Y., METASTROKE Consortium, INVENT Consortium, Sabater-Lleal, M., 2016. Genome-wide association analysis of self-reported events in 6135 individuals and 252 827 controls identifies 8 loci associated with thrombosis. Human Molecular Genetics 25, 1867–1874. doi:10.1093/hmg/ddw037.

Howrigan, D., Abbott, L., Churchhouse, C., Palmer, D., 2017. UK Biobank. http://www.nealelab.is/uk-biobank.

Jiang, F., Dong, Y., Wu, C., Yang, X., Zhao, L., Guo, J., Li, Y., Dong, J., Zheng, G.Y., Cao, H., Jin, L., Ren, Y., Cheng, W., Li, W., Tian, X.L., Li, X., 2011. Fine mapping of chromosome 3q22.3 identifies two haplotype blocks in ESYT3 associated with coronary artery disease in female Han Chinese. Atherosclerosis 218, 397–403. URL: http://www.sciencedirect.com/science/article/pii/S0021915011004989, doi:10.1016/j.atherosclerosis.2011.06.017.

Kent, W.J., Sugnet, C.W., Furey, T.S., Roskin, K.M., Pringle, T.H., Zahler, A.M., Haussler, D., 2002. The Human Genome Browser at UCSC 12, 996–1006.

Kohane, I.S., 2011. Using electronic health records to drive discovery in disease genomics. Nature Reviews Genetics 12, 417–428. URL: http://www.nature.com/articles/nrg2999, doi:10.1038/nrg2999.

Kujovich, J.L., 2011. Factor V Leiden thrombophilia. Genetics in Medicine: Official Journal of the American College of Medical Genetics 13, 1–16. doi:10.1097/GIM.0b013e3181faa0f2.

Ligthart, S., Vaez, A., Hsu, Y.H., Inflammation Working Group of the CHARGE Consortium, PMI-WG-XCP, LifeLines Cohort Study, Stolk, R., Uitterlinden, A.G., Hofman, A., Alizadeh, B.Z., Franco, O.H., Dehghan, A., 2016. Bivariate genome-wide association study identifies novel pleiotropic loci for lipids and inflammation. BMC Genomics 17, 443. URL: http://bmcgenomics.biomedcentral.com/articles/10.1186/s12864-016-2712-4, doi:10.1186/s12864-016-2712-4.

Liu, J.Z., Erlich, Y., Pickrell, J.K., 2017. Caseâcontrol association mapping by proxy using family history of disease. Nature Genetics 49, 325–331. URL: http://www.nature.com/articles/ng.3766, doi:10.1038/ng.3766.

Lucas, A.,. hudson: R package. https://github.com/anastasia-lucas/hudson.

Maher, B., 2008. Personal genomes: The case of the missing heritability. Nature 456, 18–21. doi:10.1038/456018a.

Malik, R., Chauhan, G., Traylor, M., Sargurupremraj, M., Okada, Y., Mishra, A., Rutten-Jacobs, L., Giese, A.K., van der Laan, S.W., Gretarsdottir, S., et al., 2018a. Multiancestry genome-wide association study of 520,000 subjects identifies 32 loci associated with stroke and stroke subtypes. Nature Genetics 50, 1–14. doi:10.1038/s41588-018-0058-3.

Malik, R., Rannikmae, K., Traylor, M., Georgakis, M.K., Sargurupremraj, M., Markus, H.S., Hopewell, J.C., Debette, S., Sudlow, C.L.M., Dichgans, M., for the MEGASTROKE consortium and the International Stroke Genetics Consortium, 2018b. Genome-wide meta-analysis identifies 3 novel loci associated with stroke: MEGASTROKE and UK Biobank GWAS. Annals of Neurology 84, 934–939. URL: http://doi.wiley.com/10.1002/ana.25369, doi:10.1002/ana.25369.

McCarthy, M.I., Abecasis, G.R., Cardon, L.R., Goldstein, D.B., Little, J., Ioannidis, J.P.A., Hirschhorn, J.N., 2008. Genome-wide association studies for complex traits: consensus, uncertainty and challenges. Nature Reviews Genetics 9, 356–369. URL: http://www.nature.com/articles/nrg2344, doi:10.1038/nrg2344.

National Center for Biotechnology Information, National Library of Medicine,. Database of single nucleotide polymorphisms (dbsnp). http://www.ncbi.nlm.nih.gov/SNP/. (dbSNP Build ID: GRCh37.p13), Bethesda, MD.

National Library of Medicine, 2019a. UMLS Metathesaurus - RXNORM. URL: https://www.nlm.nih.gov/research/umls/sourcereleasedocs/current/RXNORM/metadata.html.

National Library of Medicine, 2019b. UMLS SNOMED CT to ICD-10-CM Map. URL: https://www.nlm.nih.gov/research/umls/mapping\_projects/snomedct\_to\_icd10cm.html.

Neale, B., 2018. UK Biobank. URL: http://www.nealelab.is/uk-biobank.

Nielsen, J.B., Thorolfsdottir, R.B., Fritsche, L.G., Zhou, W., Skov, M.W., Graham, S.E., Herron, T.J., McCarthy, S., Schmidt, E.M., Sveinbjornsson, G., Surakka, I., Mathis, M.R., Yamazaki, M., Crawford, R.D., Gabrielsen, M.E., Skogholt, A.H., Holmen, O.L., Lin, M., Wolford, B.N., Dey, R., Dalen, H., Sulem, P., Chung, J.H., Backman, J.D., Arnar, D.O., Thorsteinsdottir, U., Baras, A., O’Dushlaine, C., Holst, A.G., Wen, X., Hornsby, W., Dewey, F.E., Boehnke, M., Kheterpal, S., Mukherjee, B., Lee, S., Kang, H.M., Holm, H., Kitzman, J., Shavit, J.A., Jalife, J., Brummett, C.M., Teslovich, T.M., Carey, D.J., Gudbjartsson, D.F., Stefansson, K., Abecasis, G.R., Hveem, K., Willer, C.J., 2018. Biobank-driven genomic discovery yields new insight into atrial fibrillation biology. Nat. Genet. 50, 1234–1239. doi:10.1038/s41588-018-0171-3.

Nikpay, M., Goel, A., Won, H.H., Hall, L.M., Willenborg, C., Kanoni, S., Saleheen, D., Kyriakou, T., Nelson, C.P., Hopewell, J.C., Webb, T.R., Zeng, L., Dehghan, A., Alver, M., Armasu, S.M., Auro, K., Bjonnes, A., Chasman, D.I., Chen, S., Ford, I., Franceschini, N., Gieger, C., Grace, C., Gustafsson, S., Huang, J., Hwang, S.J., Kim, Y.K., Kleber, M.E., Lau, K.W., Lu, X., Lu, Y., Lyytikäinen, L.P., Mihailov, E., Morrison, A.C., Pervjakova, N., Qu, L., Rose, L.M., Salfati, E., Saxena, R., Scholz, M., Smith, A.V., Tikkanen, E., Uitterlinden, A., Yang, X., Zhang, W., Zhao, W., de Andrade, M., de Vries, P.S., van Zuydam, N.R., Anand, S.S., Bertram, L., Beutner, F., Dedoussis, G., Frossard, P., Gauguier, D., Goodall, A.H., Gottesman, O., Haber, M., Han, B.G., Huang, J., Jalilzadeh, S., Kessler, T., König, I.R., Lannfelt, L., Lieb, W., Lind, L., Lindgren, C.M., Lokki, M.L., Magnusson, P.K., Mallick, N.H., Mehra, N., Meitinger, T., Memon, F.U.R., Morris, A.P., Nieminen, M.S., Pedersen, N. L., Peters, A., Rallidis, L.S., Rasheed, A., Samuel, M., Shah, S.H., Sinisalo, J., Stirrups, K.E., Trompet, S., Wang, L., Zaman, K.S., Ardissino, D., Boerwinkle, E., Borecki, I.B., Bottinger, E.P., Buring, J.E., Chambers, J.C., Collins, R., Cupples, L.A., Danesh, J., Demuth, I., Elosua, R., Epstein, S.E., Esko, T., Feitosa, M.F., Franco, O.H., Franzosi, M.G., Granger, C.B., Gu, D., Gudnason, V., Hall, A.S., Hamsten, A., Harris, T.B., Hazen, S.L., Hengstenberg, C., Hofman, A., Ingelsson, E., Iribarren, C., Jukema, J.W., Karhunen, P.J., Kim, B.J., Kooner, J.S., Kullo, I.J., Lehtimäki, T., Loos, R.J.F., Melander, O., Metspalu, A., März, W., Palmer, C.N., Perola, M., Quertermous, T., Rader, D.J., Ridker, P.M., Ripatti, S., Roberts, R., Salomaa, V., Sanghera, D.K., Schwartz, S.M., Seedorf, U., Stewart, A.F., Stott, D.J., Thiery, J., Zalloua, P.A., O’Donnell, C.J., Reilly, M.P., Assimes, T.L., Thompson, J.R., Erdmann, J., Clarke, R., Watkins, H., Kathiresan, S., McPherson, R., Deloukas, P., Schunkert, H., Samani, N.J., Farrall, M., 2015. A comprehensive 1,000 Genomes-based genome-wide association meta-analysis of coronary artery disease. Nat. Genet. 47, 1121–1130. doi:10.1038/ng.3396.

Risch, N., 1990. Linkage Strategies for Genetically Complex Traits. 1. Multilocus Models. Am J Hum Genet. 46, 222–228.

Ruderfer, D.M., Walsh, C.G., Aguirre, M.W., Tanigawa, Y., Ribeiro, J.D., Franklin, J. C., Rivas, M.A., 2019. Significant shared heritability underlies suicide attempt and clinically predicted probability of attempting suicide. Molecular Psychiatry, 1–9 URL: https://www.nature.com/articles/s41380-018-0326-8, doi:10.1038/s41380-018-0326-8.

Schnier, C., Bush, K., Nolan, J., Sudlow, C.L.M., UK Biobank Outcome Adjuducation Group, 2017. Definitions of Stroke for UK Biobank Phase 1 Outcomes Adjudication. URL: https://biobank.ctsu.ox.ac.uk/crystal/crystal/docs/alg_outcome_stroke.pdf.

Schoonjans, A.S., Meuwissen, M., Reyniers, E., Kooy, F., Ceulemans, B., 2016. PLCB1 epileptic encephalopathies; Review and expansion of the phenotypic spectrum. European journal of paediatric neurology: EJPN: official journal of the European Paediatric Neurology Society 20, 474–479. doi:10.1016/j.ejpn.2016.01.002.

Seabold, S., Perktold, J., 2010. Statsmodels: Econometric and statistical modeling with python, in: 9th Python in Science Conference.

Sinnott, J.A., Cai, F., Yu, S., Hejblum, B.P., Hong, C., Kohane, I.S., Liao, K.P., 2018. PheProb: probabilistic phenotyping using diagnosis codes to improve power for genetic association studies. Journal of the American Medical Informatics Association 25, 1359–1365. URL: https://academic.oup.com/jamia/article/25/10/1359/4999078, doi:10.1093/jamia/ocy056.

Song, Y., Ma, R., Zhang, H., 2019. The influence of MRAS gene variants on ischemic stroke and serum lipid levels in Chinese Han population. Medicine 98, e18065. URL: https://journals.lww.com/md-journal/FullText/2019/11290/The_influence_of_MRAS_gene_variants_on_ischemic.32.aspx, doi:10.1097/MD.0000000000018065.

Thangaraj, P.M., Kummer, B.R., Lorberbaum, T., Elkind, M.V.S., Tatonetti, N.P., 2019. Comparative analysis, applications, and interpretation of electronic health record-based stroke phenotyping methods. bioRxiv URL: https://www.biorxiv.org/content/early/2019/06/22/565671, doi:10.1101/565671.

The Emerging Risk Factors Collaboration, 2009. Lipoprotein(a) Concentration and the Risk of Coronary Heart Disease, Stroke, and Nonvascular Mortality. JAMA : the journal of the American Medical Association 302, 412–423. URL: https://www.ncbi.nlm.nih.gov/pmc/articles/PMC3272390/, doi:10.1001/jama.2009.1063.

Traylor, M., Tozer, D.J., Croall, I.D., Lisiecka-Ford, D.M., Olorunda, A.O., Boncoraglio, G., Dichgans, M., Lemmens, R., Rosand, J., Rost, N.S., Rothwell, P.M., Sudlow, C.L.M., Thijs, V., Rutten-Jacobs, L., Markus, H.S., International Stroke Genetics Consortium, 2019. Genetic variation in PLEKHG1 is associated with white matter hyperintensities (n = 11,226). Neurology 92, e749–e757. doi:10.1212/WNL.0000000000006952.

Turner, S.D., 2014. qqman: an R package for visualizing GWAS results using Q-Q and manhattan plots. bioRxiv, 005165 URL: https://www.biorxiv.org/content/10.1101/005165v1, doi:10.1101/005165.

UK Biobank,. Genotyping and quality control of UK Biobank, a large-scale, extensively phenotyped prospective resource. URL: http://www.ukbiobank.ac.uk/wp-content/uploads/2014/04/UKBiobank_genotyping_QC_documentation-web.pdf.

Wang Long, Chen Juan, Zeng Ying, Wei Jie, Jing Jinjin, Li Ge, Su Li, Tang Xiaojun, Wu Tangchun, Zhou Li, 2016. Functional Variant in the SLC22a3-LPAL2-LPA Gene Cluster Contributes to the Severity of Coronary Artery Disease. Arteriosclerosis, Thrombosis, and Vascular Biology 36, 1989–1996. URL: https://www.ahajournals.org/doi/full/10.1161/ATVBAHA.116.307311, doi:10.1161/ATVBAHA.116.307311.

Weiskopf, N.G., Hripcsak, G., Swaminathan, S., Weng, C., 2013. Defining and measuring completeness of electronic health records for secondary use. Journal of biomedical informatics 46, 830–836.

Wells, H.R., Freidin, M.B., Zainul Abidin, F.N., Payton, A., Dawes, P., Munro, K.J., Morton, C.C., Moore, D.R., Dawson, S.J., Williams, F.M., 2019. GWAS Identifies 44 Independent Associated Genomic Loci for Self-Reported Adult Hearing Difficulty in UK Biobank. The American Journal of Human Genetics 105, 788–802. URL: https://linkinghub.elsevier.com/retrieve/pii/S0002929719303477, doi:10.1016/j.ajhg.2019.09.008.

Whetzel, P.L., Noy, N.F., Shah, N.H., Alexander, P.R., Nyulas, C., Tudorache, T., Musen, M.A., 2011. BioPortal: enhanced functionality via new Web services from the National Center for Biomedical Ontology to access and use ontologies in software applications. Nucleic Acids Research 39, W541–W545. URL: https://academic.oup.com/nar/article-lookup/doi/10.1093/nar/gkr469, doi:10.1093/nar/gkr469.

Willer, C.J., Schmidt, E.M., Sengupta, S., Peloso, G.M., Gustafsson, S., Kanoni, S., Ganna, A., Chen, J., Buchkovich, M.L., Mora, S., Beckmann, J.S., Bragg-Gresham, J.L., Chang, H.Y., Demirkan, A., Den Hertog, H.M., Do, R., Donnelly, L.A., Ehret, G.B., Esko, T., Feitosa, M.F., Ferreira, T., Fischer, K., Fontanillas, P., Fraser, R.M., Freitag, D.F., Gurdasani, D., Heikkilä, K., Hyppönen, E., Isaacs, A., Jackson, A.U., Johansson, Å., Johnson, T., Kaakinen, M., Kettunen, J., Kleber, M.E., Li, X., Luan, J., Lyytikäinen, L.P., Magnusson, P.K.E., Mangino, M., Mihailov, E., Montasser, M.E., Müller-Nurasyid, M., Nolte, I.M., O’Connell, J.R., Palmer, C.D., Perola, M., Petersen, A.K., Sanna, S., Saxena, R., Service, S.K., Shah, S., Shungin, D., Sidore, C., Song, C., Strawbridge, R.J., Surakka, I., Tanaka, T., Teslovich, T.M., Thorleifsson, G., Van den Herik, E.G., Voight, B.F., Volcik, K.A., Waite, L. L., Wong, A., Wu, Y., Zhang, W., Absher, D., Asiki, G., Barroso, I., Been, L.F., Bolton, J.L., Bonnycastle, L.L., Brambilla, P., Burnett, M.S., Cesana, G., Dimitriou, M., Doney, A.S.F., Döring, A., Elliott, P., Epstein, S.E., Ingi Eyjolfsson, G., Gigante, B., Goodarzi, M.O., Grallert, H., Gravito, M.L., Groves, C.J., Hallmans, G., Hartikainen, A.L., Hayward, C., Hernandez, D., Hicks, A.A., Holm, H., Hung, Y.J., Illig, T., Jones, M.R., Kaleebu, P., Kastelein, J.J.P., Khaw, K.T., Kim, E., Klopp, N., Komulainen, P., Kumari, M., Langenberg, C., Lehtimäki, T., Lin, S.Y., Lindström, J., Loos, R.J.F., Mach, F., McArdle, W.L., Meisinger, C., Mitchell, B.D., Müller, G., Nagaraja, R., Narisu, N., Nieminen, T.V.M., Nsubuga, R.N., Olafsson, I., Ong, K.K., Palotie, A., Papamarkou, T., Pomilla, C., Pouta, A., Rader, D.J., Reilly, M.P., Ridker, P.M., Rivadeneira, F., Rudan, I., Ruokonen, A., Samani, N., Scharnagl, H., Seeley, J., Silander, K., Stančáková, A., Stirrups, K., Swift, A.J., Tiret, L., Uitterlinden, A.G., van Pelt, L.J., Vedantam, S., Wainwright, N., Wijmenga, C., Wild, S.H., Willemsen, G., Wilsgaard, T., Wilson, J.F., Young, E. H., Zhao, J.H., Adair, L.S., Arveiler, D., Assimes, T.L., Bandinelli, S., Bennett, F., Bochud, M., Boehm, B.O., Boomsma, D.I., Borecki, I.B., Bornstein, S.R., Bovet, P., Burnier, M., Campbell, H., Chakravarti, A., Chambers, J.C., Chen, Y.D.I., Collins, F. S., Cooper, R.S., Danesh, J., Dedoussis, G., de Faire, U., Feranil, A.B., Ferrières, J., Ferrucci, L., Freimer, N.B., Gieger, C., Groop, L.C., Gudnason, V., Gyllensten, U., Hamsten, A., Harris, T.B., Hingorani, A., Hirschhorn, J.N., Hofman, A., Hovingh, G.K., Hsiung, C.A., Humphries, S.E., Hunt, S.C., Hveem, K., Iribarren, C., Järvelin, M.R., Jula, A., Kähönen, M., Kaprio, J., Kesäniemi, A., Kivimaki, M., Kooner, J.S., Koudstaal, P.J., Krauss, R.M., Kuh, D., Kuusisto, J., Kyvik, K. O., Laakso, M., Lakka, T.A., Lind, L., Lindgren, C.M., Martin, N.G., März, W., McCarthy, M.I., McKenzie, C.A., Meneton, P., Metspalu, A., Moilanen, L., Morris, A.D., Munroe, P.B., Njølstad, I., Pedersen, N.L., Power, C., Pramstaller, P.P., Price, J.F., Psaty, B.M., Quertermous, T., Rauramaa, R., Saleheen, D., Salomaa, V., Sanghera, D.K., Saramies, J., Schwarz, P.E.H., Sheu, W.H.H., Shuldiner, A.R., Siegbahn, A., Spector, T.D., Stefansson, K., Strachan, D.P., Tayo, B.O., Tremoli, E., Tuomilehto, J., Uusitupa, M., van Duijn, C.M., Vollenweider, P., Wallentin, L., Wareham, N.J., Whitfield, J.B., Wolffenbuttel, B.H.R., Ordovas, J.M., Boerwinkle, E., Palmer, C.N.A., Thorsteinsdottir, U., Chasman, D.I., Rotter, J.I., Franks, P.W., Ripatti, S., Cupples, L.A., Sandhu, M.S., Rich, S.S., Boehnke, M., Deloukas, P., Kathiresan, S., Mohlke, K.L., Ingelsson, E., Abecasis, G.R., Global Lipids Genetics Consortium, 2013. Discovery and refinement of loci associated with lipid levels. Nat. Genet. 45, 1274–1283. doi:10.1038/ng.2797.

Xue, A., Wu, Y., Zhu, Z., Zhang, F., Kemper, K.E., Zheng, Z., Yengo, L., Lloyd-Jones, L.R., Sidorenko, J., Wu, Y., eQTLGen Consortium, McRae, A.F., Visscher, P.M., Zeng, J., Yang, J., 2018. Genome-wide association analyses identify 143 risk variants and putative regulatory mechanisms for type 2 diabetes. Nat Commun 9, 2941. doi:10.1038/s41467-018-04951-w.

Yang, J., Ferreira, T., Morris, A.P., Medland, S.E., Genetic Investigation of AN-thropometric Traits (GIANT) Consortium, DIAbetes Genetics Replication And Meta-analysis (DIAGRAM) Consortium, Madden, P.A.F., Heath, A.C., Martin, N.G., Montgomery, G.W., Weedon, M.N., Loos, R.J., Frayling, T.M., McCarthy, M.I., Hirschhorn, J.N., Goddard, M.E., Visscher, P.M., 2012. Conditional and joint multiple-SNP analysis of GWAS summary statistics identifies additional variants influencing complex traits. Nature Genetics 44, 369–375. URL: http://www.nature.com/articles/ng.2213, doi:10.1038/ng.2213.

Yang, J., Wray, N.R., Visscher, P.M., 2009. Comparing apples and oranges: equating the power of case-control and quantitative trait association studies. Genetic Epidemiology, n/a-n/a URL: http://doi.wiley.com/10.1002/gepi.20456, doi:10.1002/gepi.20456.

Zaitlen, N., Pasaniuc, B., Patterson, N., Pollack, S., Voight, B., Groop, L., Altshuler, D., Henderson, B.E., Kolonel, L.N., Marchand, L.L., Waters, K., Haiman, C.A., Stranger, B.E., Dermitzakis, E.T., Kraft, P., Price, A.L., 2012. Analysis of case-control association studies with known risk variants. Bioinformatics 28, 1729–1737. URL: https://academic.oup.com/bioinformatics/article-lookup/doi/10.1093/bioinformatics/bts259, doi:10.1093/bioinformatics/bts259.

Zaykin, D.V., Zhivotovsky, L.A., 2005. Ranks of Genuine Associations in Whole-Genome Scans. Genetics 171, 813–823. URL: http://www.genetics.org/lookup/doi/10.1534/genetics.105.044206, doi:10.1534/genetics.105.044206.

Zhang, X., Guan, J., Guo, M., Dai, H., Cai, S., Zhou, C., Wang, Y., Qin, Q., 2019. Rho GTPaseâactivating protein 1 promotes apoptosis of myocardial cells in an ischemic cardiomyopathy model. Kardiologia Polska 77, 1163–1169. doi:10.33963/KP.15040.

Zheng, J., Erzurumluoglu, A.M., Elsworth, B.L., Kemp, J.P., Howe, L., Haycock, P.C., Hemani, G., Tansey, K., Laurin, C., Pourcain, B.S., Warrington, N.M., Finucane, H.K., Price, A.L., Bulik-Sullivan, B.K., Anttila, V., Paternoster, L., Gaunt, T.R., Evans, D.M., Neale, B.M., 2017. LD Hub: a centralized database and web interface to perform LD score regression that maximizes the potential of summary level GWAS data for SNP heritability and genetic correlation analysis. Bioinformatics 33, 272–279. URL: https://academic.oup.com/bioinformatics/article/33/2/272/2525718, doi:10.1093/bioinformatics/btw613. publisher: Oxford Academic.

Zheng, P.F., Yin, R.X., Deng, G.X., Guan, Y.Z., Wei, B.L., Liu, C.X., 2019. Association between the XKR6 rs7819412 SNP and serum lipid levels and the risk of coronary artery disease and ischemic stroke. BMC Cardiovascular Disorders 19, 202. URL: https://doi.org/10.1186/s12872-019-1179-z, doi:10.1186/s12872-019-1179-z.

Zhou, W., Nielsen, J.B., Fritsche, L.G., Dey, R., Gabrielsen, M.E., Wolford, B.N., LeFaive, J., VandeHaar, P., Gagliano, S.A., Gifford, A., Bastarache, L.A., Wei, W.Q., Denny, J.C., Lin, M., Hveem, K., Kang, H.M., Abecasis, G.R., Willer, C.J., Lee, S., 2018. Efficiently controlling for case-control imbalance and sample relatedness in large-scale genetic association studies. Nature Genetics 50, 1335–1341. URL: https://www.nature.com/articles/s41588-018-0184-y, doi:10.1038/s41588-018-0184-y.

